# Convergent relaxation of molecular constraint in mammalian herbivores highlights the roles of liver and kidney functions in carnivory

**DOI:** 10.1101/2023.11.17.567625

**Authors:** Matthew D. Pollard, Wynn K. Meyer, Emily E. Puckett

## Abstract

Mammalia comprises a great diversity of diet types and associated adaptations. An understanding of the genomic mechanisms underlying these adaptations may offer insights for improving human health. Comparative genomic studies of diet that employ taxonomically restricted analyses or simplified diet classifications may suffer reduced power to detect molecular convergence associated with diet evolution. Here, we used a quantitative carnivory score—indicative of the amount of animal protein in the diet—for 80 mammalian species to detect significant correlations between the relative evolutionary rates of genes and changes in diet. We identified six genes—*ACADSB*, *CLDN16*, *CPB1*, *PNLIP*, *SLC13A2*, and *SLC14A2*—that experienced significant changes in evolutionary constraint alongside changes in carnivory score, becoming less constrained in lineages evolving more herbivorous diets. We further considered the biological functions associated with diet evolution and observed that pathways related to amino acid and lipid metabolism, biological oxidation, and small molecule transport experienced reduced purifying selection as lineages became more herbivorous. Liver and kidney functions showed similar patterns of constraint with dietary change. Our results indicate that, in highly carnivorous lineages, selection acts on the liver and kidneys to maintain sufficient metabolism and excretion of substances found in excess in carnivorous diets. These biological functions become less important with the evolution of increasing herbivory, so experience a relaxation of constraint in more herbivorous lineages.

## INTRODUCTION

As a diversity of mammalian diets arose from the ancestral insectivorous strategy (Gill et al. 2014), numerous physiological, morphological, and behavioral adaptations also evolved. Understanding the mechanisms of these adaptations may lead to improvements in human health. For example, glucose metabolism in healthy carnivores resembles diabetes in humans and other non-carnivores (Schermerhorn 2013). Polar bears are adapted to persistently high blood cholesterol levels (Liu et al. 2014), which have been implicated in human cardiovascular disease (Ference et al. 2017), and some networks underlying polar bear adaptation have been implicated in high fat diet adaptation in humans (Fumagalli et al. 2015). Thus, comparative genomic studies of diet offer novel insights for medical advancement. Large-scale comparative genomic resources, consisting of data from hundreds of non-model mammals, have been developed to support such research (Zoonomia Consortium 2020; Christmas et al. 2023).

Previous studies identified many genomic changes accompanying adaptations to carnivorous and herbivorous diets. For example, the convergent evolution of herbivory in the giant and red pandas coincided with adaptive molecular evolution of genes associated with the utilization of nutrients that are scarce in bamboo, as well as limb development genes that facilitated growth of the pseudothumb (Hu et al. 2017). In cetaceans, positive selection for proteinases and lipases and loss of pancreatic *RNASE1* expression have been attributed to the evolution of carnivory from a herbivorous ancestor (Wang et al. 2016). Loss of the hormone-receptor pair *INSL5*-*RXFP4*, which regulates appetite and glucose homeostasis, is thought to be an adaptation to irregular feeding patterns in carnivores (Hecker et al. 2019). Positive selection in carnivores acted on genes associated with successful hunting—such as muscle strength and agility—as well as digestion of fat-and protein-rich foods (Kim et al. 2016).

While previous studies provided valuable insights into lineage-specific dietary adaptations, studies that incorporate data encompassing a larger number of taxa can identify mechanisms shared across convergent changes in diet. To date, comparative genomic studies that sampled broadly within mammals have used coarse dietary classification systems, such as herbivore, carnivore, and omnivore (Kim et al. 2016; Hecker et al. 2019; Wu 2022). This results in a loss of important dietary information, as species grouped together differ in the proportions of plant and animal matter consumed and may show divergent adaptations (Pineda-Munoz and Alroy 2014; Grundler and Rabosky 2020; Pollard and Puckett 2022; Reuter et al. 2023). Thus, the power of comparative genomic studies that investigate the molecular mechanisms of dietary adaptation may be reduced.

The objectives of our study are two-fold. First, we identify genes that are important for convergent adaptation to increasingly carnivorous or herbivorous diets in mammals. We accomplish this by detecting significant correlations between gene evolutionary rates and changes in a continuous diet score mapped across a phylogeny of 80 species. Second, we use convergent evolutionary rate shifts to elucidate the biological functions associated with adaptation to increasingly carnivorous or herbivorous diets across Mammalia. Our analyses encompass unprecedented taxonomic breadth and dietary nuance compared to previous comparative genomic studies of diet in mammals.

## RESULTS

### Molecular evolutionary rates associated with dietary change

We generated a carnivory score for each species from the proportion of animal matter in their diet, as reported in the EltonTraits dataset (Wilman et al. 2014). Thus, our study treats invertivorous species as carnivorous. A score of 0 represents a completely herbivorous species, and a score of 100 a completely carnivorous mammal. Due to the structure of EltonTraits, carnivory scores jumped in steps of 10 from 0 to 100 (Fig. 1; Supplemental Table S1); we therefore analyzed 11 ordinal bins hereafter called degrees of carnivory.

**Figure 1.**
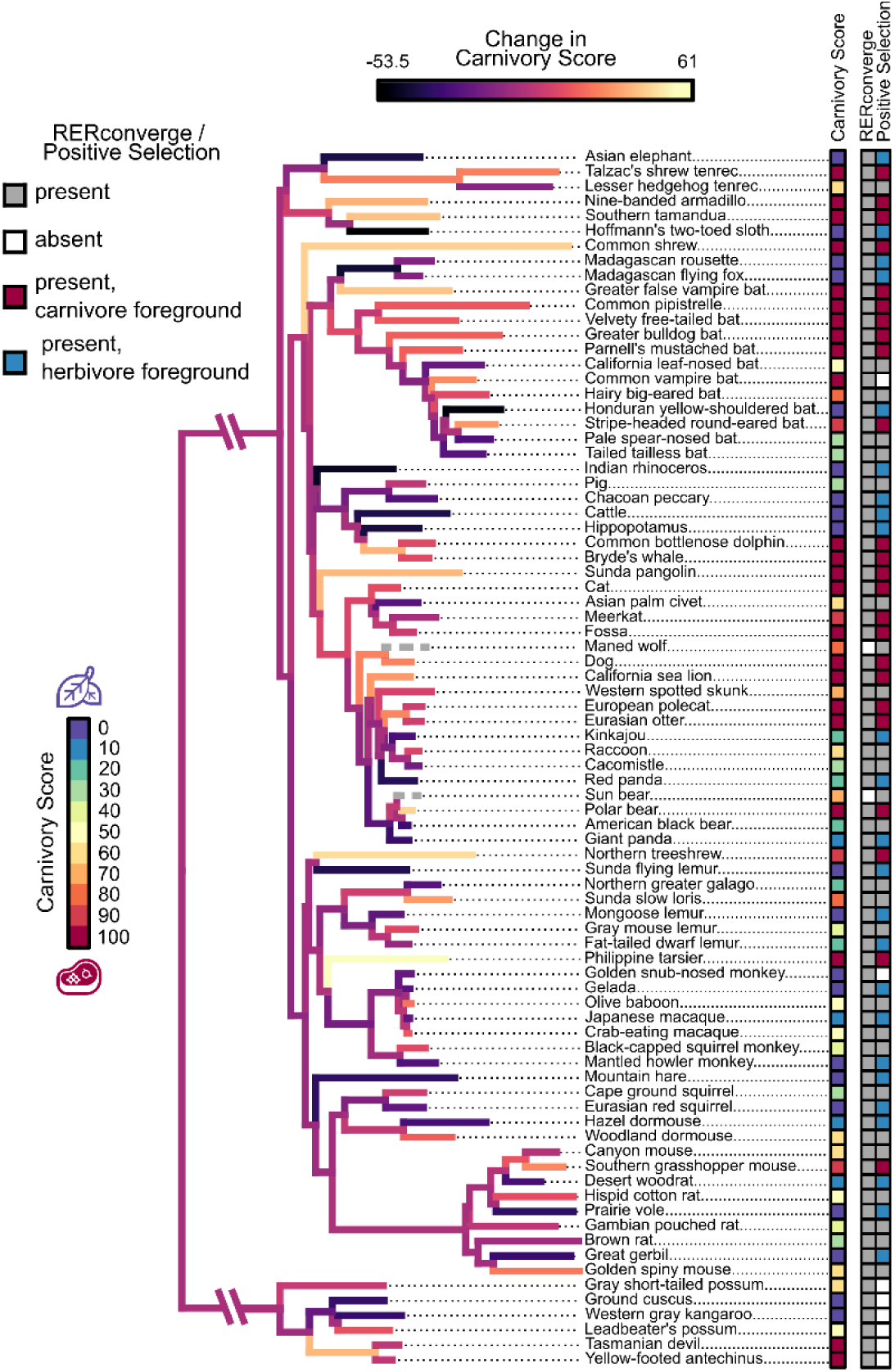
Species selection for comparative genomic analyses of mammalian diet. The carnivory score represents the proportion of the diet composed of animal-based food items for each species, as listed in EltonTraits (Wilman et al. 2014). This score was used as input in our RERconverge analyses and informed selection of foreground species in our tests for positive selection. Branch colors represent the magnitude and direction of change in carnivory score across the phylogeny, as inferred by using fast estimation of maximum likelihood ancestral states (Revell 2012). Bright yellow and dark purple branches indicate increases and decreases in carnivory score, respectively. Gray dashed branches represent species—maned wolf and sun bear—that were not included in our RERconverge analyses and so were not used to estimate change in carnivory score. Branch length represents the average evolutionary rate across all genes for a given branch of the maximum clade credibility phylogeny of Upham et al. (2019), as reported by RERconverge. Most species were included in both the RERconverge and positive selection analyses and are marked as present (gray, red, blue) in the corresponding columns. Species not included in an analysis are marked as absent (white). For the positive selection analyses, species included in the carnivorous (carnivory score ≥90) and herbivorous (carnivory score ≤10) foregrounds are marked in the corresponding column as red and blue, respectively. Scientific names and carnivory scores are provided for each species in Supplemental Table S1.

We identified genes that experienced convergent evolutionary constraint or positive selection in association with changes in carnivory across a phylogeny of 80 mammal species (Fig. 1). Of 13,912 genes included in our RERconverge analysis (Kowalczyk et al. 2019), we identified six with a significant negative correlation between relative evolutionary rate (RER) and change in carnivory score (FDR=0.05; Table 1; Fig. 2A; Supplemental Table S3). These significant correlations are a result of convergent rate changes in multiple regions of the phylogeny (Supplemental Fig. S1). A negative correlation represents the following pattern: the greater the decrease in carnivory, the higher the RER of the gene. Decreasing carnivory is also proportional to increasing herbivory. Strong negative correlations may be driven by increasingly rapid evolution as the change in carnivory score becomes increasingly negative, slower evolution as the change in carnivory score becomes increasingly positive, or both. Hereafter, we refer to genes with this pattern of association as "negatively correlated with change in carnivory score". A positive correlation represents the opposite pattern: the greater the increase in carnivory (equivalent to decreasing herbivory), the higher the relative rate of evolution of the gene.

**Figure 2.**
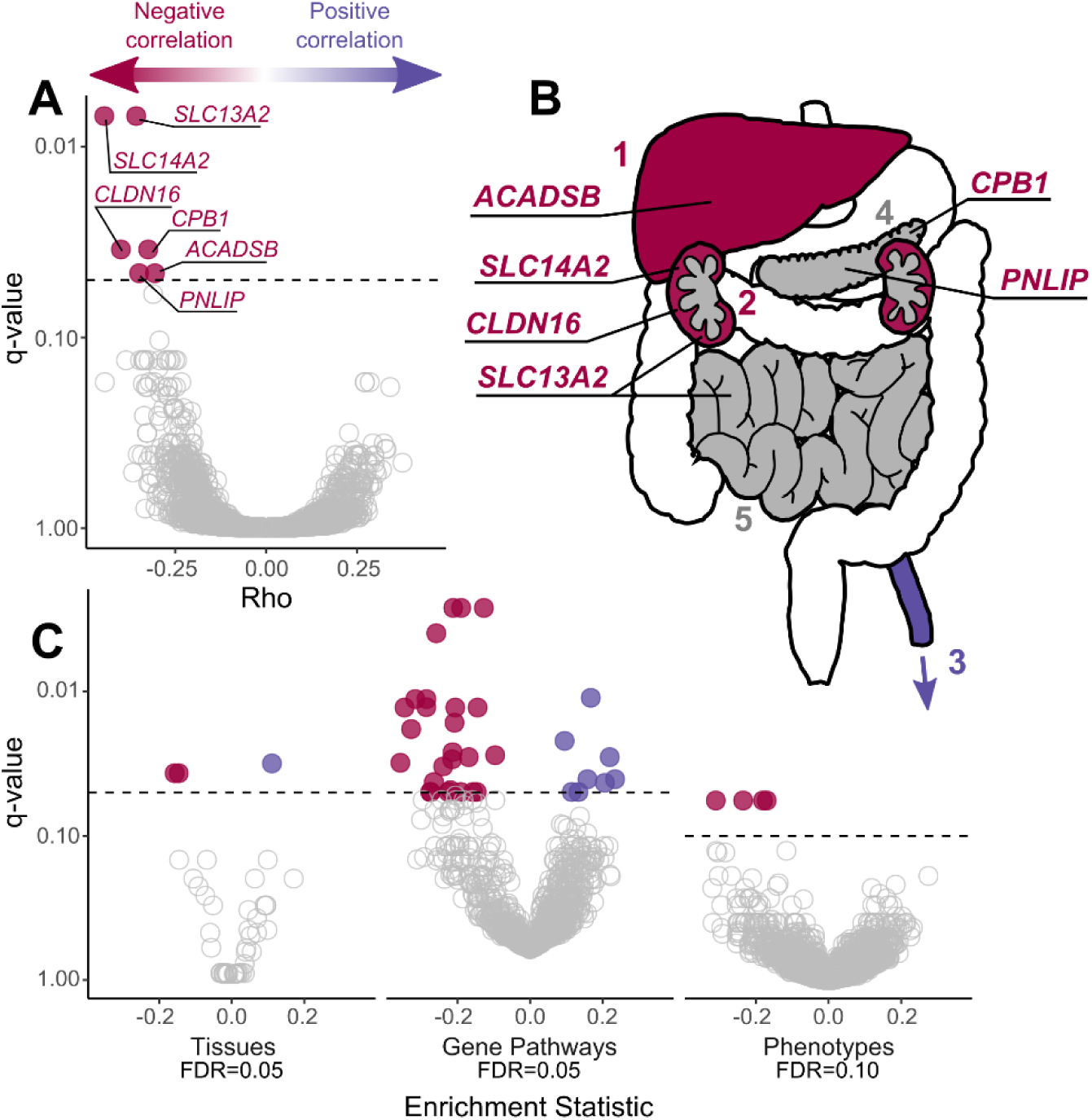
Top genes and pathways with signatures of diet-associated evolutionary constraint. (*A*) Genes identified by RERconverge as having a significant association between relative evolutionary rate and change in carnivory score. A negative correlation (Rho) signifies the following pattern: the greater the decrease in carnivory, the higher the rate of evolution of the gene. A positive correlation indicates the opposite pattern. After 100,000 permulations, six genes showed a significant association, and each evolved faster in association with decreased carnivory (FDR=0.05). (*B*) Tissues enriched for positive (purple) and negative (red) correlations, where: 1 = liver, 2 = kidneys, 3 = tibial artery, 4 = pancreas, 5 = small intestine. Some tissues are annotated with the genes found to be individually significant. The annotated tissues are the locations of highest expression for those genes in adult humans, according to the Genotype-Tissue Expression Project (2015). Gray represents organs that were not significantly enriched but were sites of strongest expression individually significant genes. (*C*) Biological functions showing constraint associated with change in diet. We used RERconverge to test for enrichment of gene sets representing tissues (n=50), gene pathways (n=1,290), and abnormal phenotypes (n=3,560). For tissues, two gene sets were enriched for negative correlations and one set was enriched for positive correlations (Supplemental Table S4). For gene pathways, 26 gene sets were enriched for negative correlations and 8 sets were enriched for positive correlations (Supplemental Table S5). For abnormal phenotypes, five gene sets were enriched for negative correlations (Supplemental Table S6).

**Table 1.** Genes with evolutionary rates that are significantly associated with change in carnivory score. Permulation *P*-values represent the proportion of 100,000 permulations that produced a stronger correlation with change in carnivory score than the observed value for each gene. Multiple hypothesis testing corrections were performed by generating *Q*-values using Storey’s correction method (Storey et al. 2020; FDR=0.05). A negative correlation (Rho) signifies the following pattern: the greater the decrease in carnivory, the higher the rate of evolution of the gene.

| Gene | Rho | Parametric<br>*P*-value | Parametric<br>*Q*-value | Permutation<br>*P*-value | Permutation<br>*Q*-value |
| --- | --- | --- | --- | --- | --- |
| *SLC14A2* | -0.446 | $1.54 \times 10^{-8}$ | $2.14 \times 10^{-4}$ | $<1.00 \times 10^{-5}$* | $<0.035$* |
| *SLC13A2* | -0.358 | $6.94 \times 10^{-6}$ | 0.037 | $<1.00 \times 10^{-5}$* | $<0.035$* |
| *CLDN16* | -0.400 | $7.93 \times 10^{-6}$ | 0.037 | $1.00 \times 10^{-5}$ | 0.035 |
| *CPB1* | -0.325 | $8.02 \times 10^{-5}$ | 0.150 | $1.00 \times 10^{-5}$ | 0.035 |
| *PNLIP* | -0.350 | $1.85 \times 10^{-5}$ | 0.065 | $2.00 \times 10^{-5}$ | 0.046 |
| *ACADSB* | -0.306 | $3.36 \times 10^{-4}$ | 0.260 | $2.00 \times 10^{-5}$ | 0.046 |

\* After generating 100,000 null statistics, none produced a stronger correlation with diet than the observed values for *SLC13A2* and *SLC14A2*. However, if the *P*-values are adjusted to the smallest observed non-zero *P*-value ( $1.00 \times 10^{-5}$ ), they would produce a significant empirical *Q*-value (FDR=0.05).

Negatively correlated genes make a greater contribution to fitness the more carnivory score increases, or a lower contribution the more carnivory score decreases (Kowalczyk et al. 2020). Among genes with this signature, the products of three (*SLC13A2*, *SLC14A2*, and *CLDN16*) are involved in transport or resorption in the kidney, two (*CPB1* and *PNLIP*) are secreted pancreatic enzymes, and one (*ACADSB*) is involved in amino acid metabolism (Davis et al. 1991; Pajor 1999; Simon et al. 1999; Andresen et al. 2000; Leung and Morser 2018; Geng et al. 2020). Our analysis had greater power to detect negative associations between RERs and change in carnivory score compared to positive associations. The results of a Mann-Whitney *U* test indicated a significant difference in the distribution of empirical, permulation-based *P*-values for genes with positive and negative correlations (AUC=0.523, *P*=2.16 x 10^-6^), with fewer low *P*-values generated for positively correlated genes (Supplemental Fig. S2).

Each of the six significant genes are most strongly expressed in the liver, kidneys, pancreas, or small intestine in humans (The GTEx Consortium 2015). We performed a gene set enrichment analysis to determine if there was a statistically significant association between change in carnivory score and the rate of evolution of genes expressed in specific tissues (n = 50). Our test showed that the liver and kidney cortex gene sets were significantly enriched for genes that are negatively correlated with change in carnivory score (FDR=0.05; Fig. 2B,C; Tables 2, S4). In contrast, the tibial artery was significantly enriched for genes evolving more quickly the more carnivory score increased.

**Table 2.** Tissues that are significantly enriched for genes associated with change in carnivory score. Permulation *P*-values represent the proportion of 100,000 permulations that produced a stronger enrichment test statistic than the observed value for each gene set. Multiple hypothesis testing corrections were performed by generating *Q*-values using Storey’s correction method (Storey et al. 2020; FDR=0.05). A negative enrichment statistic signifies the following pattern: the greater the decrease in carnivory, the higher the rate of evolution of the gene.

| Tissue | Enrichment Statistic | Parametric $P$-value | Parametric $Q$-value | Permutation $P$-value | Permutation $Q$-value |
| --- | --- | --- | --- | --- | --- |
| Tibial Artery | 0.111 | $2.64 \times 10^{-4}$ | 0.002 | $6.30 \times 10^{-4}$ | 0.032 |
| Liver | -0.158 | $1.28 \times 10^{-19}$ | $6.38 \times 10^{-18}$ | 0.002 | 0.037 |
| Kidney Cortex | -0.146 | $7.42 \times 10^{-7}$ | $1.24 \times 10^{-5}$ | 0.002 | 0.037 |

We extended our enrichment analysis to 1,290 gene pathways from the Reactome and KEGG databases and identified significant enrichment for 34 pathways (FDR=0.05; Fig. 2C; Supplemental Table S5). Of these pathways, 26 were associated with faster evolution the more carnivory score decreases. These pathways had functions related to digestion, biological oxidation, transport of small molecules, heme degradation, and metabolism of amino acids, lipids, and carbohydrates (Fig. 3). Eight pathways evolved at faster rates the more carnivory score increases. These pathways were associated with cardiovascular disease, cell motility, signaling by anaplastic lymphoma kinase (ALK) in cancer, and signaling by Hippo, NOTCH3, and interleukin-17 (Supplemental Fig. S3). Our gene correlation and pathway enrichment results were robust to changes in species choice and ancestral reconstruction methodology (Supplemental Figs. S4-S5).

**Figure 3.**
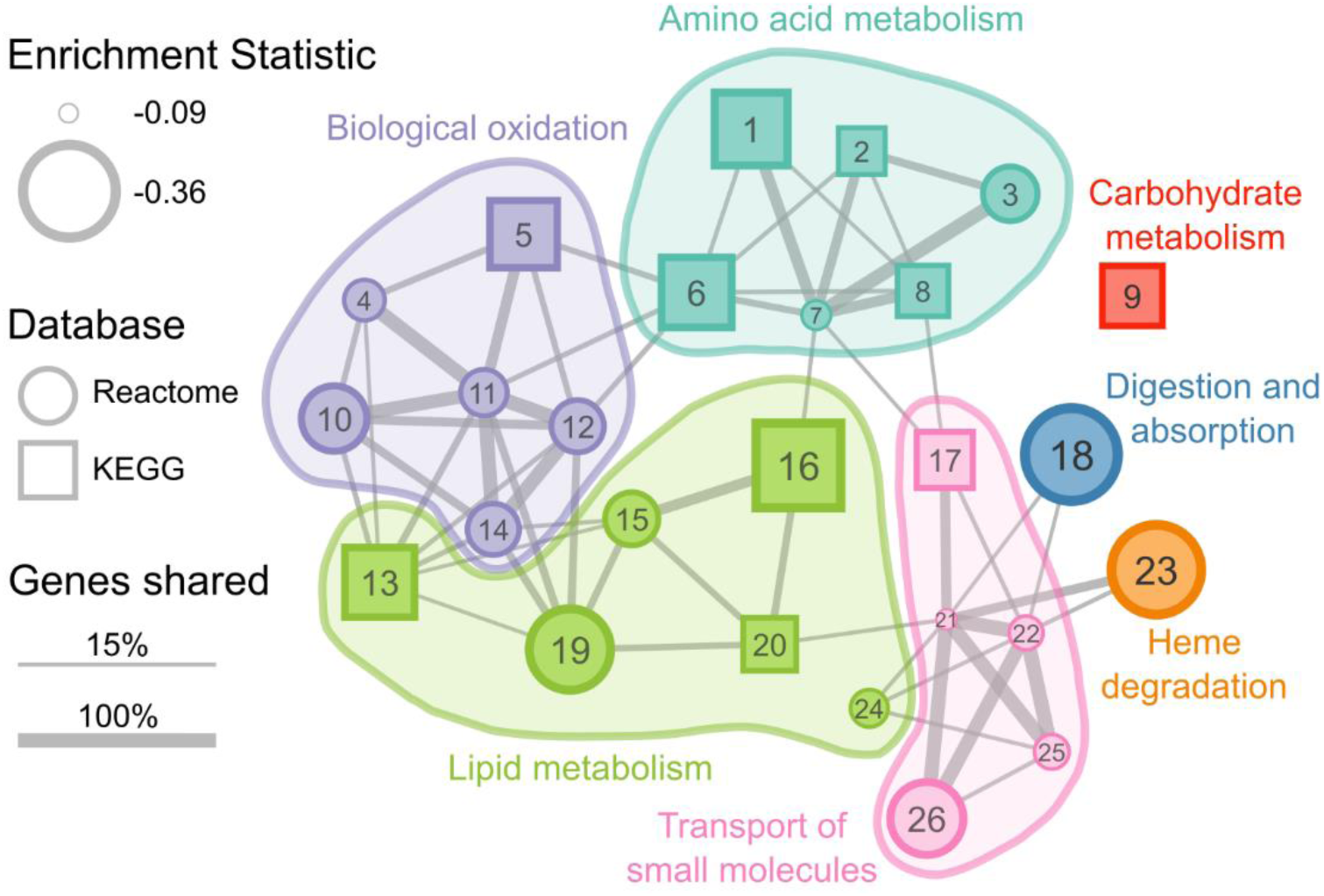
Significantly enriched gene pathways (n=26) under less evolutionary constraint as carnivory score decreases. Each circle or square represents a gene pathway. Circles and squares represent pathways from the Reactome and KEGG databases, respectively. The size of the shape represents the magnitude of the difference between the distribution of test statistics for genes in that pathway and the distribution for all other genes. Larger shapes, representing pathways with more negative correlation statistics, indicate greater reductions in evolutionary constraint as carnivory score decreases. The width of lines connecting pathways represents the proportion of shared genes in the smaller gene set. Colors represent the broad functional categories that the pathways occupy. 1, Glycine, serine, and threonine metabolism; 2, Cysteine and methionine metabolism; 3, Sulfur amino acid metabolism; 4, Phase II – Conjugation of compounds; 5, Drug metabolism – Cytochrome P450; 6, Phenylalanine metabolism; 7, Metabolism of amino acids and derivatives; 8, Arginine and proline metabolism; 9, Starch and sucrose metabolism; 10, Metabolic disorders of biological oxidation enzymes; 11, Biological oxidations; 12, Phase I – Functionalization of compounds; 13, Steroid hormone biosynthesis; 14, Cytochrome P450 – Arranged by substrate type; 15, Fatty acid metabolism (Reactome); 16, Fatty acid metabolism (KEGG); 17, Proximal tubule bicarbonate reclamation; 18, Digestion and absorption; 19, Synthesis of bile acids and bile salts via 7alpha-hydroxycholesterol; 20, PPAR signaling pathway; 21, Transport of small molecules; 22, SLC-mediated transmembrane transport; 23, Heme degradation; 24, Synthesis of phosphatidylcholine; 25, Transport of bile salts and organic acids, metal ions, and amine compounds; 26, Amino acid transport across the plasma membrane. A table of results for all gene pathways tested (n=1,290) is available in Supplemental Table S5.

We performed an enrichment analysis on 3,560 gene sets representing abnormal phenotypes from the Mammalian Phenotype Ontology (Smith et al. 2005). We identified five phenotypes with significant enrichment using both raw and permulation-based empirical *P*-values (FDR=0.1; Fig. 2C; Table 3; Supplemental Table S6). All five gene sets were enriched for genes evolving faster as carnivory score decreases. Two phenotypes—decreased urine osmolality (MP:0002988) and increased urine calcium level (MP:0005441)—were related to abnormal urine homeostasis, and two others—abnormal lipid level (MGI:0001547) and abnormal circulating lipid level (MGI:0003949)—were associated with lipid homeostasis. The abnormal urine homeostasis gene sets shared 8 of 82 total genes, with 4—*SLC12A1*, *KCTD1*, *SLC4A1*, and *UMOD*—in the 10 top-ranked genes for both gene sets. The abnormal lipid homeostasis gene sets shared 10 of 134 total genes, with 1 overlapping gene, *BHMT*, in the top 10 for both sets.

**Table 3.** Phenotype-based gene sets from the Mammalian Phenotype Ontology that are significantly nriched for genes associated with carnivory score. Permulation *P*-values represent the proportion of 100,000 ermulations that produced a stronger enrichment statistic than the observed value for each gene set. Multiple hypothesis testing corrections were performed by generating *Q*-values using Storey’s correction method (Storey t al. 2020; FDR=0.1). A negative enrichment statistic signifies the following pattern: the greater the decrease in arnivory, the higher the rate of evolution of the gene.

| Phenotype | Enrichment Statistic | Parametric *P*-value | Parametric *Q*-value | Permutation *P*-value | Permutation *Q*-value |
| --- | --- | --- | --- | --- | --- |
| MP:0002223 lymphoid hypoplasia | -0.304 | $8.60 \times 10^{-4}$ | 0.076 | $<7.00 \times 10^{-5*}$ | $<0.057*$ |
| MP:0003949 abnormal circulating lipid level | -0.235 | $3.30 \times 10^{-5}$ | 0.011 | $7.00 \times 10^{-5*}$ | 0.057* |
| MP:0001547 abnormal lipid level | -0.170 | $3.62 \times 10^{-9}$ | $1.20 \times 10^{-5}$ | $8.00 \times 10^{-5}$ | 0.057 |
| MP:0005441 increased urine calcium level | -0.311 | $7.74 \times 10^{-8}$ | $8.53 \times 10^{-5}$ | $8.00 \times 10^{-5}$ | 0.057 |
| MP:0002988 decreased urine osmolality | -0.181 | $1.01 \times 10^{-5}$ | 0.006 | $8.00 \times 10^{-5}$ | 0.057 |

\* After generating 100,000 null statistics, none produced a stronger correlation with diet than the observed values for MP:0002223. However, if the *P*-values are adjusted to the smallest observed non-zero *P*-value ( $7.00 \times 10^{-5}$ ), they would produce a significant empirical *Q*-value (FDR=0.1).

### Molecular evolutionary rates associated with binary diet classifications

For the significant genes that we identified in our continuous RERconverge analysis, the negative correlations between RER and change in carnivory score appeared to be most strongly driven by large RERs coinciding with the greatest decreases in score (i.e., shifts towards increasing herbivory). Separately calculating Pearson’s correlation coefficients for increases and decreases in carnivory score supported this, as significant relationships with RER were only identified for decreases in carnivory score (Fig. 4). To determine if this translated to separate relationships with RER at opposing ends of the carnivory score spectrum, we ran binary RER analyses with either the most carnivorous (carnivory score ≥90) or most herbivorous (carnivory score ≤10) lineages selected as the foreground. Using hypercarnivorous lineages as the foreground, we observed that no genes showed a significant difference in RER between foreground and background lineages (FDR=0.05; Supplemental Table S7). The same was true for the most herbivorous lineages (FDR=0.05), however we did identify 28 genes with marginally significant associations (FDR<0.1) in this analysis (Supplemental Table S8). This included four of the six genes identified in our analysis of continuous carnivory score—*ACADSB*, *CPB1*, *PNLIP*, and *SLC13A2*—although the strengths of the associations were greater in the continuous analysis. We identified significant enrichment for 78 KEGG and Reactome pathways in the herbivore-specific analysis (FDR=0.05), and 21 of these overlapped with the 34 pathways enriched in the continuous analysis (Supplemental Table S9). Pearson’s correlation coefficients and linear regression analyses indicated that the resulting test statistics of the hyperherbivore analysis were more strongly correlated with those of the continuous analysis than were the test statistics of the hypercarnivore analysis (Supplemental Figs. S6-7).

**Figure 4.**
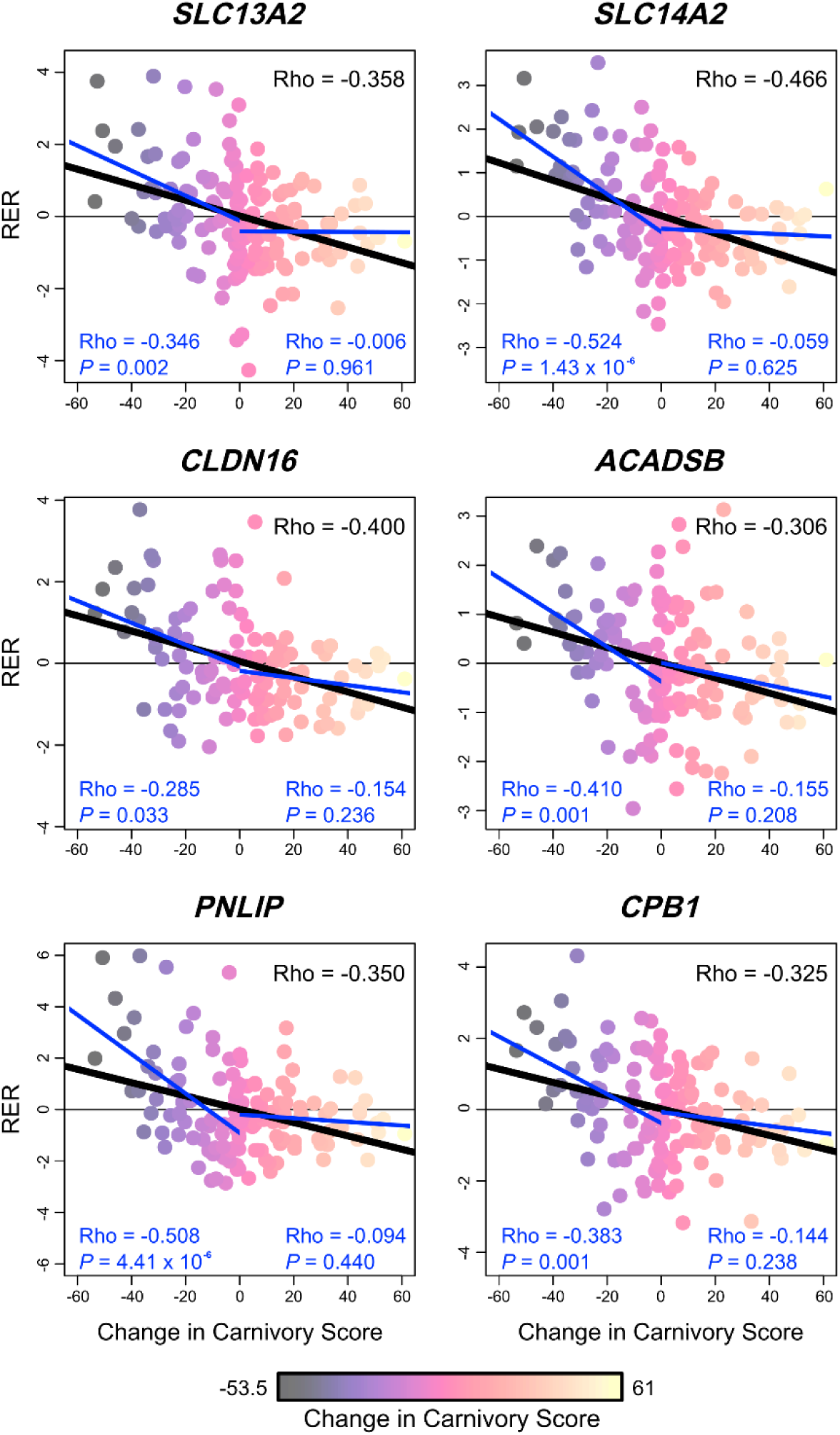
Relative evolutionary rates (RERs) associated with changes in carnivory score across Mammalia for six diet-associated genes: *SLC13A2*, *SLC14A2*, *CLDN16*, *ACADSB*, *PNLIP*, and *CPB1*. As in Fig. 1, light yellow and darker purple points indicate increases and decreases in carnivory score, respectively. Lines of best fit represent the relationship between RER and change in carnivory score as inferred by linear regression. Pearson’s correlation coefficients (Rho) represent the strength and direction of correlations (upper right values) between RER and change in carnivory score. Black lines of best fit and correlation coefficients represent the relationship across all changes in carnivory score, as inferred by our continuous RERconverge analysis. Blue lines, with associated correlations, represent the relationship with RER for only decreases (left) or increases (right) in carnivory score.

### Positive selection

We identified 193 and 172 genes that showed significant evidence of positive selection in the most carnivorous and herbivorous species, respectively (FDR=0.05; Supplemental Tables S10, S11). Among the carnivore-specific positively selected genes, we identified enrichment for four Reactome pathways related to *O*-linked glycosylation and one pathway related to pregnenolone biosynthesis (Table 4). The *ADAMTS* (a disintegrin and metalloproteinase with thrombospondin motifs) genes driving the observed enrichment for *O*-linked glycosylation pathways were also components of the only GO Term that was significantly enriched—metallopeptidase activity (GO:0008237; Table 4). We did not identify significant enrichment of any phenotypes, gene pathways, or GO Terms in our list of positively selected genes in herbivores (FDR=0.05).

**Table 4.** Gene sets enriched for positive selection in the most carnivorous mammals (carnivory score ≥90). We did not detect enrichment in the most herbivorous mammals (carnivory score ≤10).

| Gene Set | # Genes analyzed | Positively selected genes | P-value | Q-value |
| --- | --- | --- | --- | --- |
| Reactome Pathways |  |  |  |  |
| *O*-linked glycosylation | 87 | *ADAMTS5*, *ADAMTS12*,<br>*ADAMTS14*, *ADAMTS20*,<br>*B4GALT6*, *CHST4*, *LARGE2*,<br>*THSD4* | $1.83 \times 10^{-5}$ | 0.008 |
| Defective *B3GLCT* causes Peters-plus syndrome (PpS) | 34 | *ADAMTS5*, *ADAMTS12*,<br>*ADAMTS14*, *ADAMTS20*,<br>*THSD4* | $7.70 \times 10^{-5}$ | 0.013 |
| *O*-glycosylation of TSR domain-containing proteins | 35 | *ADAMTS5*, *ADAMTS12*,<br>*ADAMTS14*, *ADAMTS20*,<br>*THSD4* | $8.89 \times 10^{-5}$ | 0.013 |
| Diseases associated with *O*-glycosylation of proteins | 49 | *ADAMTS5*, *ADAMTS12*,<br>*ADAMTS14*, *ADAMTS20*,<br>*THSD4* | $4.50 \times 10^{-4}$ | 0.041 |
| Pregnenolone biosynthesis | 12 | *CYP11A1*, *FDX2*, *STARD3NL* | $4.58 \times 10^{-4}$ | 0.041 |
| GO Terms |  |  |  |  |
| GO:0008237 metallopeptidase activity | 158 | *ADAMTS5*, *ADAMTS12*,<br>*ADAMTS14*, *ADAMTS20*,<br>*AEBP1*, *CNDP1*, *CPZ*, *PEPD*,<br>*STAMBPL1*, *THSD4*, *TRHDE* | $5.75 \times 10^{-6}$ | 0.002 |

## DISCUSSION

We used quantitative carnivory scores and a phylogeny of 80 species sampled from the taxonomic and dietary breadth of Mammalia to identify genes that vary in functional importance based on the relative proportion of animal-based food sources in a diet. Ours is the first comparative genomic study of diet to use a quantitative variable and to analyze convergence across so many species. In our analyses of relative evolutionary rates, we were better able to identify signals of convergent relaxation as carnivory decreases than the opposite pattern. Our analyses of quantitative carnivory score changes are complementary to but distinct from methods that search for the most important genes in the most carnivorous or herbivorous lineages (i.e., without considering all increases and decreases in carnivory score, as in our binary analyses of RER), and better account for the continuous nature of this complex trait. The main targets of convergence in our study were lipid and amino acid metabolism, biological oxidation, and liver and kidney functions (Fig. 2; Tables 2-4, S5, S6).

Many of the signals observed in our analyses appear to be driven most strongly by rapid evolution in increasingly herbivorous lineages (Fig. 4). Indeed, a binary analysis of RERs revealed that many of the biological functions enriched in the continuous analyses were similarly enriched among genes evolving rapidly in hyperherbivores. However, the inverse is not true for carnivorous lineages (Supplemental Tables S7-S9). While increasingly herbivorous diets mediated much of the signal detected in our analyses, the negatively correlated genes and functions are of greater relevance to carnivory than herbivory. The faster evolution detected during increases in herbivory represents a convergent relaxation—and decrease in importance—of genes and functions that are otherwise crucial to carnivory. Rapid evolution due to positive selection is transient and less likely to be detected by RERconverge than sustained relaxation of purifying selection (Kowalczyk et al. 2020). This is supported by the lack of overlap between our RERconverge and positive selection results (Supplemental Tables S3, S7-8, S10-11).

### The roles of liver and kidneys in carnivory

Multiple lines of evidence support that liver and kidney functions are crucial for carnivory in mammals. Of the six genes identified by our continuous RERconverge analysis as having a significant negative correlation between RER and change in carnivory score, three (*SLC13A2*, *SLC14A2*, and *CLDN16*) and one (*ACADSB*) are respectively most strongly expressed in human kidneys and liver (The GTEx Consortium 2015). Further, the kidney cortex and liver gene sets were enriched for negatively correlated genes (Table 2).

The importance of kidney function to carnivory may be mediated by its role in the reabsorption and secretion of small molecules. Our enrichment analysis identified several pathways related to small molecule transport, including several containing solute carrier proteins (SLCs) such as the products of *SLC13A2* and *SLC14A2*, that were enriched for genes with negative correlations between RER and change in carnivory score. Many of these pathways take place in the kidneys. The results of our phenotype-specific enrichment analysis also support a central role of small molecule transportation in the kidneys of lineages with increasingly carnivorous diets. Of the five MGI phenotype gene sets that were significantly enriched, two of them—decreased urine osmolality (MP:0002988) and increased urine calcium level (MP:0005441)—are related to abnormal urine homeostasis, which is a direct consequence of impaired renal function. Mammals excrete nitrogenous waste in the form of urea, and the urea load that must be excreted each day largely depends on the protein content of an organism’s diet. The most carnivorous species have the greatest urea loads and produce urine with higher urea concentrations than other species (Liu et al. 2011). The need to excrete greater quantities of nitrogenous waste imposes selective pressure on the carnivore kidney. This is supported by our finding that the evolutionary rate of *SLC14A2* increases as carnivory score decreases, as the urea transporters encoded by this gene are critical to the urea-dependent urine concentrating mechanism (Geng et al. 2020).

Liver function is an important component of adaptation to carnivory due to its role in amino acid and lipid metabolism. It is the only organ in mammals that can completely metabolize most amino acids, and it plays an active role in amino acid synthesis (Hou et al. 2020). Carnivorous diets are rich in proteins and lipids. Pathways related to their metabolism should therefore be under stronger purifying selection the more carnivory increases, as lipids and amino acids represent carnivores’ main energy sources. Among negatively correlated gene pathways, we identified enrichment for genes involved in several processes related to the metabolism of lipids and assorted amino acids, including many that take place within the liver. One such pathway was related to bile acid synthesis (pathway 19, Fig. 3). Bile acids facilitate digestion of dietary fats and allow excess cholesterol to be excreted from the body. Indeed, bile acid synthesis and excretion represent the major mechanisms of cholesterol catabolism and elimination in mammals (Russell 2003). Given that carnivores possess lipid-rich diets, lipid metabolism pathways may be evolutionarily constrained so that sufficient levels of cholesterol homeostasis are maintained in these species. Lipid homeostasis may be less important in lineages with lipid-poor diets, so constraint relaxes as increasingly herbivorous lineages evolve.

Metabolism of amino acids and lipids is associated with biological oxidation, and we found that several biological oxidation pathways were enriched for genes negatively correlated with change in carnivory score. Thus, biological oxidation is an integral component of carnivory. However, this direction of association is an unexpected result, given the importance of xenobiotic metabolism in herbivores. Many plants employ chemical defenses against herbivory (Berenbaum 1995), and herbivores may overcome these defenses via oxidation (Karban and Agrawal 2002; Dearing et al. 2005). Indeed, Hecker et al. (2019) found that carnivores have lost function of several genes associated with xenobiotic detoxification, as this process is less important in carnivores compared to herbivores. The negative correlation between the RERs of biological oxidation genes and change in carnivory score may be driven by rapid evolution in increasingly herbivorous lineages. In support of this, our binary RER analysis indicated that biological oxidation pathways evolve significantly faster in the most herbivorous lineages (Supplemental Table S9). However, we did not detect herbivore-specific positive selection acting on the genes that drove the enrichment of these pathways (Supplemental Table S11). This suggests that positive selection is not driving the observed rapid evolution of these genes in herbivores. Our results hint at a previously unobserved role for biological oxidation in species with more carnivorous diets.

Amino acid and lipid metabolism outside of the liver are also associated with change in carnivory. The evolutionary rates of *CPB1* and *PNLIP*—two genes most strongly expressed in the pancreas (The GTEx Consortium 2015)—were significantly negatively correlated with changing carnivory score. A gene for another pancreatic lipase, *PNLIPRP2*, had a marginally significant association with change in carnivory score (Rho=-0.311; *Q*=0.059) and may be more important in lineages where carnivory is not decreasing. The products of *PNLIP* and *PNLIPRP2* are secreted from the pancreas into the small intestine, where they enable efficient digestion of dietary fats (Davis et al. 1991; Lowe 2002). They have been implicated in the dietary switch to carnivory during the evolution of Cetacea, when lipids became a major nutritional component of whales’ diets (Wang et al. 2016; Wu 2022). Our results support previous findings that pancreatic lipases play a role during evolutionary changes in degree of carnivory. Interestingly, another member of this family of pancreatic lipase genes, *PNLIPRP1*, has convergently lost function in multiple herbivorous mammals, suggesting a relaxation of selective constraint in herbivores compared to carnivores (Hecker et al. 2019). The protein encoded by this gene, PL-RP1, shows little to no detectable lipase activity and instead acts as a competitive inhibitor of pancreatic lipase (Lowe 2002). Inactivation of *PNLIPRP1* would therefore enhance fat digestion capacity in herbivores. One may expect that inactivation of *PNLIPRP1* in herbivores may drive a significant negative correlation between RER and change in carnivory score, as this represents an extreme relaxation of selection in herbivores. However, we observed no significant association for *PNLIPRP1*, likely because the gene alignments used in our study do not include pseudogenes. Thus, relaxation of selective pressure leading to pseudogenization cannot be detected using our dataset, and this may have reduced the number of significant associations between RERs and change in carnivory score that we could observe.

Biological oxidation outside the liver is important for dietary adaptation: our positive selection analyses showed that a member of the cytochrome P450 gene superfamily, *CYP11A1*, is positively selected in the most carnivorous species. The protein encoded by *CYP11A1* catalyzes the first stage of steroid hormone synthesis—the production of pregnenolone from cholesterol via oxidation (Payne and Hales 2004). The pregnenolone biosynthesis pathway was enriched in our carnivore-specific positive selection analysis (Table 4), and steroid hormone synthesis showed less constraint as carnivory score decreases (Supplemental Table S5).

In summary, our results indicate that, when mammals increase in carnivory or maintain a high carnivory score, purifying selection acts on functions related to metabolism and elimination of substances found in excess in animal-based foods. The liver and kidneys have been disproportionately targeted to maintain sufficient functioning of these processes. Meanwhile, these functions are less important in herbivorous diets, so they experience a substantial relaxation of evolutionary constraint in lineages adapting to increasing herbivory. This relaxation drove much of the signal detected by our RERconverge analyses.

### Implications of carnivorous adaptations for human health

Our results offer insights into diet-related diseases in humans and suggest avenues for new medical research. Human dietary maladaptation is not driven by high rates of carnivory *per se*: stable isotopes and archeological evidence indicate that increased meat consumption began early in human evolution (Sponheimer and Lee-Thorp 1999; Domínguez-Rodrigo et al. 2005) and that pre-agricultural diets were more carnivorous than current ones (Eaton and Cordain 1997). Instead, an evolutionarily rapid shift away from pre-agricultural diets in favor of fattier meats and processed foods has increased chronic disease in modern communities (Jew et al. 2009). Our findings help characterize mechanisms of human dietary maladaptation because metabolic challenges posed by extreme carnivory overlap with those posed by modern processed diets. For example, carnivore adaptations that allow for lipid-rich diets without disease are medically relevant given a global increase in the consumption of high-fat foods in human populations (Kearney 2010). In humans, high intakes of red meat are associated with increased blood cholesterol levels and coronary heart disease (Al-Shaar et al. 2020). Tightly regulated cholesterol and lipid homeostasis may be an adaptation to avoid such diseases in carnivores. Our phenotype-specific enrichment analysis supports that lipid homeostasis is functionally important in lineages that maintain or increase carnivory. Two overlapping significantly enriched phenotype gene sets—abnormal lipid level (MGI:0001547) and abnormal circulating lipid level (MGI:0003949)—are related to lipid homeostasis and were enriched for genes negatively correlated with carnivory score. Thus, the lipid metabolism genes driving our results may be important targets for future study.

Evolutionary constraint on the carnivore kidney is relevant to human health due to the elevated burden of high-protein diets on renal function. High-protein diets have been linked to kidney damage and reduced renal function in observational human studies (Ko et al. 2020). In other omnivorous mammals, experimental evidence has shown that high protein intake causes kidney inflammation and damage (Jia et al. 2010; Tovar-Palacio et al. 2011). Protein-rich diets have been implicated in kidney stone formation as they increase the excretion of calcium and oxalate into the urine (Robertson et al. 1979). The link between diet and kidney stones offers a plausible connection between *SLC13A2*, carnivory, and human health, as defects in this gene cause low urinary citrate concentrations, leading to the formation of kidney stones (Okamoto et al. 2007). Defects in another of the significantly negatively correlated genes, *CLDN16*, are associated with elevated urine calcium levels and deposition of calcium salts in the kidneys (Konrad et al. 2008).

Another pathway with genes that show evidence of relaxation as carnivory score decreases is heme degradation. While nonheme iron is present in both plant-and animal-based foods, heme iron is mainly found in animal tissues (Hurrell and Egli 2010). Primarily carnivorous species ingest excess heme, so heme degradation is a more important process in these species than in omnivores or herbivores. Excess heme intake causes oxidative damage, inflammation, and cell death (Chiabrando et al. 2014), and has been implicated in several diseases, including type-2 diabetes, coronary heart disease, gut dysbiosis, colitis, and cancers (Bastide et al. 2011; Hooda et al. 2014; Constante et al. 2017). Cancer risk is highest among carnivorous mammals (Vincze et al. 2021), and selection for DNA repair mechanisms in carnivores has been attributed to increased consumption of heme-related reactive oxygen species (Kim et al. 2016). Decreased heme-related disease risk may drive the observed pattern of relaxed evolutionary constraint as carnivory score decreases.

Carnivore-specific positive selection acting on *O*-linked glycosylation pathways and metalloprotease activity may have relevance to human diabetes. Enrichment for these pathways was primarily driven by four *ADAMTS* genes. *O*-linked glycosylation is a widespread post-translational modification of ADAMTS metalloproteases (Wang et al. 2007). The positively selected ADAMTSs are associated with extracellular matrix (ECM) assembly and degradation in various biological processes (Glasson et al. 2005; McCulloch et al. 2009; El Hour et al. 2010; Dupuis et al. 2011; Dancevic et al. 2012; Dubail and Apte 2015). In mice, high-fat diets and insulin resistance increase expression of ECM remodeling genes and encourage collagen deposition, leading to fibrosis in tissues such as skeletal muscle, liver, and adipose tissue (Choi et al. 2015; Pincu et al. 2015; Williams et al. 2015). High-fat diets and healthy glucose metabolism that resembles diabetes (Schermerhorn 2013) may increase selective pressure acting on ECM-regulating proteases in carnivores.

### Comprehensive sampling elucidated fundamentally important functions

Our analyses sampled the taxonomic and dietary breadth of Mammalia, incorporating genome-wide data from 80 species. The signals of convergent molecular evolution that we detected are driven by multiple instances of dietary convergence across the phylogeny (Fig. 1; Supplemental Fig. S1). We therefore isolated genes and biological functions that are important for carnivory and herbivory in most or all instances that such diets evolved across mammals. Some of our results overlap with those previously found in studies of fewer species. For example, lipid metabolism has frequently been identified as a target of selection in carnivores (Liu et al. 2014; Kim et al. 2016; Wang et al. 2016). However, our study is the first to emphasize the fundamental importance of kidney and liver function to carnivory across mammals.

By including more species in our analyses, we provide a clearer picture of which diet-specific selection signatures are clade-specific and which are apparent in all carnivores. For example, Kim et al. (2016) found carnivore-specific selection acting on functions such as neuron development, muscle strength, and agility. The authors attribute their findings to agile hunting behaviors. However, we did not detect this selection in our broader analysis. Some carnivores in our expanded sampling are not agile predators and are unlikely to have experienced selection on these functions. For example, Bryde’s whale was included in our analyses and feeds on zooplankton with limited movement capabilities (Izadi et al. 2022). Bryde’s whale does not require the same agility while hunting as the carnivorans, orca, and Tasmanian devil that were analyzed by Kim et al. (2016).

### Increases in herbivory led to convergent relaxation more often than constraint

The evolutionary history of mammalian diet offers a potential explanation for why many of our results were driven by molecular changes that co-occurred with evolution towards herbivory. The ancestral mammalian diet was insectivorous (Gill et al. 2014) and would be treated as completely carnivorous in our study. Thus, less carnivorous diets represent derived states that are likely associated with great molecular and phenotypic change as entirely new dietary adaptations arise. In contrast, lineages that increase in carnivory are returning to a state more like the ancestral condition, potentially leading to less dramatic shifts in gene evolutionary rates. This may lead to a stronger relationship between gene evolutionary rates and diet for decreases in carnivory score.

While the history of mammalian diet may help explain the stronger relationship with gene evolutionary rates as carnivory score decreases, it does not explain why the signal of evolutionary constraint was much weaker than relaxation in this direction of dietary change (Supplemental Fig S2, Supplemental Tables S3-6, S8-9). The weaker signal may be a consequence of the base level of constraint that acts on protein evolution (Worth et al. 2009). Given that proteins already experience some level of constraint, it is likely harder to detect further reductions in an already low evolutionary rate compared to increases.

The weaker signal we observed in genes with positive correlations than in those with negative correlations could also indicate that constraint on coding sequence is less predictable during adaptation to increased herbivory than to increased carnivory. Low predictability may be mediated by diverse digestive strategies in herbivores, and by greater variation in the nutrient composition of plant-based foodstuffs. Herbivores can be categorized by the method of microbial fermentation employed to digest plant material. Foregut and hindgut fermenters differ in the volume and nutritional quality of plant matter that they can consume (Alexander 1993), and this places different morphophysiological constraints on each strategy (Clauss et al. 2003). Fruits, seeds, and foliage differ in the proportions of protein, lipids, and structural versus non-structural carbohydrates that they contain (Jordano 2000). Thus, the diets of frugivores, granivores, and folivores differ in digestibility and energy content, and each strategy will employ a different suite of adaptations to reflect this. This diversity within herbivory may help explain our observation of less convergent constraint acting during evolution of increasing herbivory compared to carnivory.

### Pathways enriched for positive correlated genes show unclear relevance to diet evolution

The relevance to diet or health of pathways containing genes that are positively correlated with change in carnivory score is generally less obvious than those with the opposite pattern. These pathways may represent understudied components of dietary adaptation. For example, Hippo signaling integrates signals from multiple sources to regulate cell proliferation and differentiation, and can influence and be influenced by the metabolism of glucose, lipids, and amino acids (Ibar and Irvine 2020). Given that multiple metabolic cues are integrated into Hippo signaling, it is unclear why genes in this pathway experience less constraint the more carnivory increases. NOTCH3 plays an important role during vascular development and in continued vascular functioning of adult organisms (Domenga et al. 2004; Loerakker et al. 2018; Hosseini-Alghaderi and Baron 2020). Given this, enrichment for NOTCH3 signaling may be related to enrichment of the tibial artery gene set in the same direction (Table 2). However, it is unclear why vascular function would be under less constraint as carnivory increases. ALK activation is involved in the development of many cancers (Della Corte et al. 2018), but its function beyond cancer is not as well understood. It is thought to play a role in nervous system development (Palmer et al. 2009), but expression has also been detected in the small intestine and colon (Morris et al. 1994). More recently, ALK signaling was implicated in regulating energy expenditure and weight gain (Orthofer et al. 2020), so it might play an important, unidentified role in diet regulation, and further study could elucidate its relevance.

Despite limited signal and unclear relevance for many pathways enriched for positive correlated genes, one pathway stood out as having a clear connection to diet evolution. The importance of interleukin-17 (IL-17) signaling in lineages adapting to more herbivorous diets may reflect IL-17’s role in gut microbiome regulation. IL-17 encourages the production of antimicrobial proteins and the migration of neutrophils into infected intestinal mucosa, maintains intestinal barrier integrity, and limits gut dysbiosis (Aujila et al. 2007; Ishigame et al. 2009; Cao et al. 2012; Pérez et al. 2018). Gut microbiomes are essential for herbivores. The plants consumed by herbivores comprise complex polysaccharides that cannot be digested by mammalian enzymes. However, microorganisms in the herbivore gut can ferment these compounds to produce metabolites that are used easily by their host. The importance of microorganisms for plant consumption has led to convergence in gut microbiota across mammalian herbivores, while those of carnivores are highly variable (Muegge et al. 2011; Zoelzer et al. 2021). As gut microbes are necessary for successful herbivory, evolutionary constraint should act on gut homeostasis pathways in lineages with decreases in carnivory score or that maintain herbivorous diets.

### Limitations of the study and suggestions for future work

Although RERconverge has demonstrated success in associating gene RERs with the evolution of continuous traits (Kowalczyk et al. 2020), the underlying model may have some limitations. When analyzing a continuous trait, RERconverge does not by default associate raw trait values with gene RERs because this would lead to phylogenetic dependence among branches. Instead, RERconverge calculates trait change between a descendent and its ancestor (Kowalczyk et al. 2019). It detects genes with RERs that are significantly correlated with trait change, as such genes are crucial to the convergent evolution of that trait. However, in lineages that remain stationary in trait space following an ancestral transition, trait-associated genes may remain under continued evolutionary constraint or relaxation, despite no change in the trait. Consequently, the model may fail to implicate these genes as important and may instead identify genes that are important only for trait transitions and not in maintaining a phenotype once it has evolved. By quantifying the robustness of our results to species removal (Supplemental Fig. S4), we demonstrated that the signals we detected were not impacted by specific lineages, regardless of whether they represent areas of the phylogeny with active or stationary diet evolution. Thus, we can be confident that the genes we identified play important roles in adaptation to changes in carnivory score across Mammalia. However, we may have had insufficient power to detect additional diet-associated genes.

We also showed congruence between our continuous and binary analyses of RER (Supplemental Tables S3, S5, S8, S9). When implemented with lineages subsequent to transitions as foreground, binary RERconverge identified genes involved in transition to and maintenance of a foreground trait. However, this is at the cost of ignoring small changes in the trait that may be evolutionarily relevant. The ability to integrate changes in dietary traits may explain why our continuous analyses returned significant associations at an FDR of 0.05 while the binary analyses did not (Supplemental Tables S3, S7-8). For future studies of continuous traits, it may be preferable to implement a version of RERconverge that calculates changes in RER, rather than correlating trait change with raw RER values. This would negate the issue caused by lineages with extreme yet unchanging trait values, because important genes would show little change in RER on the corresponding branches.

Although our consideration of degree of carnivory as a quantitative trait expands upon prior studies that define diet categorically, this definition has limitations as well. EltonTraits, used to construct our carnivory score, was built primarily from qualitative summarizations of existing literature written in Walker’s Mammals of the World (Nowak 1999). While our carnivory score is likely accurate for extreme carnivores and herbivores, accuracy of intermediate scores may vary based on how reliably the diets were coded within EltonTraits. Inaccurate diet classifications for species with intermediate phenotypes would introduce noise into the inference of how carnivory score evolves on the phylogeny, ultimately reducing the power of our RERconverge analyses. Nevertheless, EltonTraits data has been used effectively to address several questions in ecology and evolution, such as the influence of diet on the evolution of mammalian gut microbiomes (Groussin et al. 2017), the effect of diet on species’ responses to climate change (Buckley et al. 2018), and the decline of ecological and functional diversity over time (Cooke et al. 2019; Brodie et al. 2021). The demonstrated efficacy of EltonTraits in prior works suggests that it can provide valuable data for our study, despite these limitations.

Our analyses may be impacted by a mismatch between carnivory score and genomic data for the dog, *Canis lupus*. The EltonTraits entry for *C. lupus* represents the hypercarnivorous wolf, while the genetic data was obtained from a domesticated dog. During early domestication, the dog adapted to a more omnivorous diet (Axelsson et al. 2013). A carnivory score of 100 does not accurately reflect the diet of modern dogs. However, this mismatch should not strongly impact our results. Dogs certainly possess some adaptations to omnivory, but their domestication occurred relatively recently (∼15,000 years; Savolainen et al. 2002; Pang et al. 2009). Thus, signals of selection for carnivory may be detectable in dogs, given the longer evolutionary history of hypercarnivory within the lineage. Regardless of whether there is a mismatch, it is unlikely that any significant signal will be singularly mediated by the branch leading to dog.

Our quantitative carnivory score more accurately reflects the biologically continuous nature of diet and leads to less information loss compared to typical categorical classifications, but it still does not capture all dietary variation. While the diets of strictly herbivorous species may be identical in that they do not include animal matter, they differ in nutritional composition based on the plant-based foodstuffs consumed—seeds, fruit, leaves, or other plant matter. The same is true of carnivorous species. For example, the blood-based diet of the common vampire bat differs from other carnivores included in our study by being vitamin-and lipid-poor (Mendoza et al. 2018). Studies that separate nutritional composition from the carnivory-herbivory axis of diet variation may detect signals of selection that would otherwise be masked, and can test the hypotheses of earlier comparative genomic analyses of diet. For example, a recent study considered fat intake and found evidence to support an earlier hypothesis that functional loss of *PNLIPRP1* in herbivores is associated with low fat consumption (Hecker et al. 2019; Wagner et al. 2022). Insights from comparative genomic studies of diet composition will produce important insights for medical advancement in the future.

## METHODS

### Preparation of dietary data

To create a continuous categorization of diet, we obtained data from the EltonTraits mammal dataset (Wilman et al. 2014). Data compiled in EltonTraits came primarily from Walker’s Mammals of the World (Nowak 1999), which describes species’ diets based on summaries of existing literature. Qualitative descriptions of dietary preferences were translated by the EltonTraits authors into numerical values representing the proportion of diet comprising numerous food types. In this study, we generated a carnivory score for each species by summing the proportions of vertebrates and invertebrates in their diet.

### Study species and gene alignments

For our analyses of relative evolutionary rates (RERs), we selected 80 species that encompass the taxonomic and dietary breadth of therian mammals (Fig. 1; Supplemental Table S1). We filtered multiple codon alignments that were generated with TOGA and MACSE v2 by the Zoonomia Consortium (2020; Ranwez et al. 2018; Kirilenko et al. 2023) using the methodology described in Wirthlin et al. (2024; see Supplemental Methods). We then pruned the alignments so that they included only our chosen species, and removed sequences that did not cover 50% of the total aligned sequence length. For alignments that included more than one sequence per species, we retained the sequence with the highest percent identity to human. We only analyzed alignments with sequences that survived filtering for at least 60 target species. Our filtering reduced the number of genes analyzed from 17,439 to 15,117. An overrepresentation analysis of Gene Ontology (GO) terms indicated that the unanalyzed genes were predominantly associated with DNA assembly and organization, perception of chemical stimuli, and gene silencing (Supplemental Table S2). These nucleotide alignments were translated to amino acid alignments for use with RERconverge.

### Associating relative evolutionary rates with the evolution of diet

We used RERconverge in R (Kowalczyk et al. 2019) to identify genes with consistent changes in evolutionary rate associated with the evolution of the quantitative carnivory score. RERs reveal deviation in evolutionary rate along a specific branch of the phylogeny, based on genome-wide expectations. A highly positive RER may indicate positive selection or relaxation of constraint. A highly negative RER indicates that increased constraint has led to fewer substitutions than expected. As we implemented it, RERconverge estimates the correlation between RER and change in a continuous phenotype across a phylogeny. A significant negative correlation between RER and change in carnivory score corresponds to slower evolution the more carnivory increases, and/or faster evolution the more carnivory decreases. Increased evolutionary constraint is associated with slower molecular evolution and implies that the gene contributes more to fitness under a particular phenotypic condition, so we looked for significant negative correlations when identifying genes with greater functional importance during evolution of increasing carnivory (Kowalczyk et al. 2020). Conversely, for genes critical to increasing herbivory, we looked for significant positive correlations.

We used the *estimatePhangornTreesAll* wrapper function from RERconverge, which utilizes a maximum likelihood approach implemented in phangorn (Schliep 2011), to calculate branch lengths for each gene while fixing the topology of the gene trees to match the maximum clade credibility phylogeny of Upham et al. (2019). These trees were used as input for the RERconverge analyses alongside our carnivory scores. We winsorized the RERs of each gene so that the three most extreme values at each end were set to that of the next most extreme RER. This minimized the influence of outliers. We implemented the default branch length filtering in RERconverge, setting the shortest 5% of branches across all gene trees to N/A. This resulted in an additional 1,205 genes being filtered for having fewer than 60 branches remaining in the tree, leaving 13,912 remaining genes in this analysis.

To evaluate confidence in gene-trait associations inferred by RERconverge, we used a phylogenetically restricted permutation strategy (‘permulations’; Kowalczyk et al. 2020; Saputra et al. 2021). We randomly simulated carnivory scores for each species 100,000 times using a Brownian motion model of evolution. Then for each simulated set of values, the observed carnivory scores were assigned to species based on the ranks of the simulated scores. We used these shuffled phenotype sets to calculate gene-trait associations using RERconverge, generating 100,000 null statistics per gene. We calculated a new, empirical *P*-value for each gene-trait association based on the proportion of null statistics that are equally or more extreme than the observed statistic for that association. We corrected for multiple hypothesis testing using Storey’s correction method (Storey et al. 2020; FDR=0.05). Using the Mann-Whitney *U* test for enrichment available in RERconverge, we identified biological functions associated with the evolution of diet. We constructed tissue-specific gene sets from the Genotype-Tissue Expression Project (GTEx; The GTEx Consortium 2015) data, defining tissue-specificity following the strategy of Jain and Tuteja (2019). Briefly, tissue-specific genes were identified as those with expression levels of at least one transcript per million (TPM) that were significantly higher (at least five-fold) in up to seven tissues compared to all others. We tested for tissue-specific enrichment using these gene sets, gene pathway enrichment using KEGG and Reactome pathways from MSigDB (Liberzon et al. 2011), and enrichment for abnormal phenotypes using gene sets from the Mammalian Phenotype (MP) Ontology (Smith et al. 2005).

We checked the robustness of our RERconverge results to species choice and ancestral reconstruction method (see Supplemental Methods). We also compared our results to those produced using binary classifications of diet (see Supplemental Methods).

### Positive selection associated with the evolution of carnivory or herbivory

We tested each gene for positive selection via branch-site models (branch-site neutral versus branch-site selection) using codeml from PAML (Yang 2007), as implemented in BASE (Forni et al. 2021). We also used BUSTED from the HyPhy software suite (Murrell et al. 2015; Pond et al. 2020). These models test for positive selection in a group of foreground species chosen *a priori*. Terminal branches leading to species with carnivory scores ≥90 (n=25) and ≤10 (n=24) were selected as foreground for carnivory-and herbivory-specific analyses, respectively (Fig. 1; Supplemental Table S1). Select species with carnivory scores of 20—specifically the red panda and kinkajou—were included in the herbivore foreground despite relatively high carnivory scores because they are considered in the broader literature to be primarily herbivorous (Yonzon and Hunter 1991; Julien-Laferrière 1999; Kays 1999; Pradhan et al. 2001; Panthi et al. 2012).

As in our RERconverge analyses, we obtained nucleotide alignments for each gene from Zoonomia and applied the same filtering steps. However, due to an updated data release from Zoonomia, the alignment files used for positive selection tests differed from those obtained for the RERconverge analysis. Specifically, the second set included the maned wolf and sun bear, but did not include marsupials. Given that the maned wolf and sun bear have carnivory scores that differ from their closely related species (Fig. 1), we retained them in these analyses to increase our representation of dietary diversity across the phylogeny.

PAML branch-site models do not test for the absence of positive selection in background species, so these tests alone do not identify genes experiencing positive selection in association with a trait of interest (Kowalczyk et al. 2021). To confirm that our significant branch-site results corresponded to foreground-specific positive selection, we removed the foreground species and tested for tree-wide positive selection using site models (M1a versus M2a) in PAML. Only genes detected as positively selected by the branch-site model and not the site model with foreground species excluded were retained (Supplemental Fig. S8).

We employed a conservative strategy to generate final lists of positively selected genes. To account for the impact of gene tree discordance on our results (Mendes and Hahn 2016), we performed positive selection analyses using gene trees and the fixed species tree. We used RAxML v8.2.12 (Stamatakis 2014) with the GTRGAMMAI nucleotide substitution model to generate gene trees. We only considered a gene to have experienced diet-associated positive selection when it produced a significant result in both our codeml and BUSTED analyses, and if this result persisted when using the gene tree and the species tree (Supplemental Fig. S8). As an additional filtering step, we checked that the signals of positive selection returned by codeml and BUSTED were not caused by relaxation of selection by using RELAX (Wertheim et al. 2015), and excluded any genes showing significant relaxation from our final gene lists (Supplemental Fig. S8). We used overrepresentation analyses to test our final lists for enrichment of GO terms, gene pathways from the Reactome and KEGG databases, phenotypes from the MP Ontology, and our tissue-specific gene sets using clusterProfiler and ReactomePA (Yu et al. 2012; Yu and He 2016). For all overrepresentation analyses, we used human annotation databases.

## DATA ACCESS

All nucleotide and amino acid alignments used in this study were generated via TOGA annotation of orthologous genes by the Zoonomia Consortium (https://zoonomiaproject.org/the-data/; https://genome.senckenberg.de/download/TOGA/human_hg38_reference/MultipleCodonAlignments/). The filtered nucleotide and amino acid alignments generated in this study are available on FigShare (https://figshare.com/s/133b5237f20e82986312; https://doi.org/10.6084/m9.figshare.25695762). Carnivory scores for species included in this study are available in Supplemental Table S1. All RERconverge and positive selection results generated in this study are available in Supplemental Tables S3-11. All code used for data processing and analysis are available on GitHub (https://github.com/mdpllard/carnivory-genes), FigShare, and as Supplemental Code. The tissue-and mouse phenotype-specific gene sets generated in this study are available on GitHub, FigShare, and in the Supplemental Code.

## ACKNOWLEDGEMENTS

We thank the administrators of the University of Memphis High Performance Computing (HPC) facilities for support during this study. We thank the three anonymous reviewers for their insightful feedback on the submitted manuscript, Amanda Kowalczyk and Ruby Redlich for developing the ‘getPermsBinaryFudged’ function to improve permulation speed, and Emily Kopania and Michael Tene for helpful discussions. This material is based upon work supported by the National Science Foundation under Grant No. 2233124 to WKM.

## AUTHOR CONTRIBUTIONS

MDP and WKM performed data pre-processing. MDP analyzed the data and interpreted results with EEP and WKM. MDP wrote the manuscript. EEP and WKM edited the manuscript. EEP supervised the study.

