## Supplemental_Figures for "Convergent relaxation of molecular constraint in mammalian herbivores highlights the roles of liver and kidney functions in carnivory"

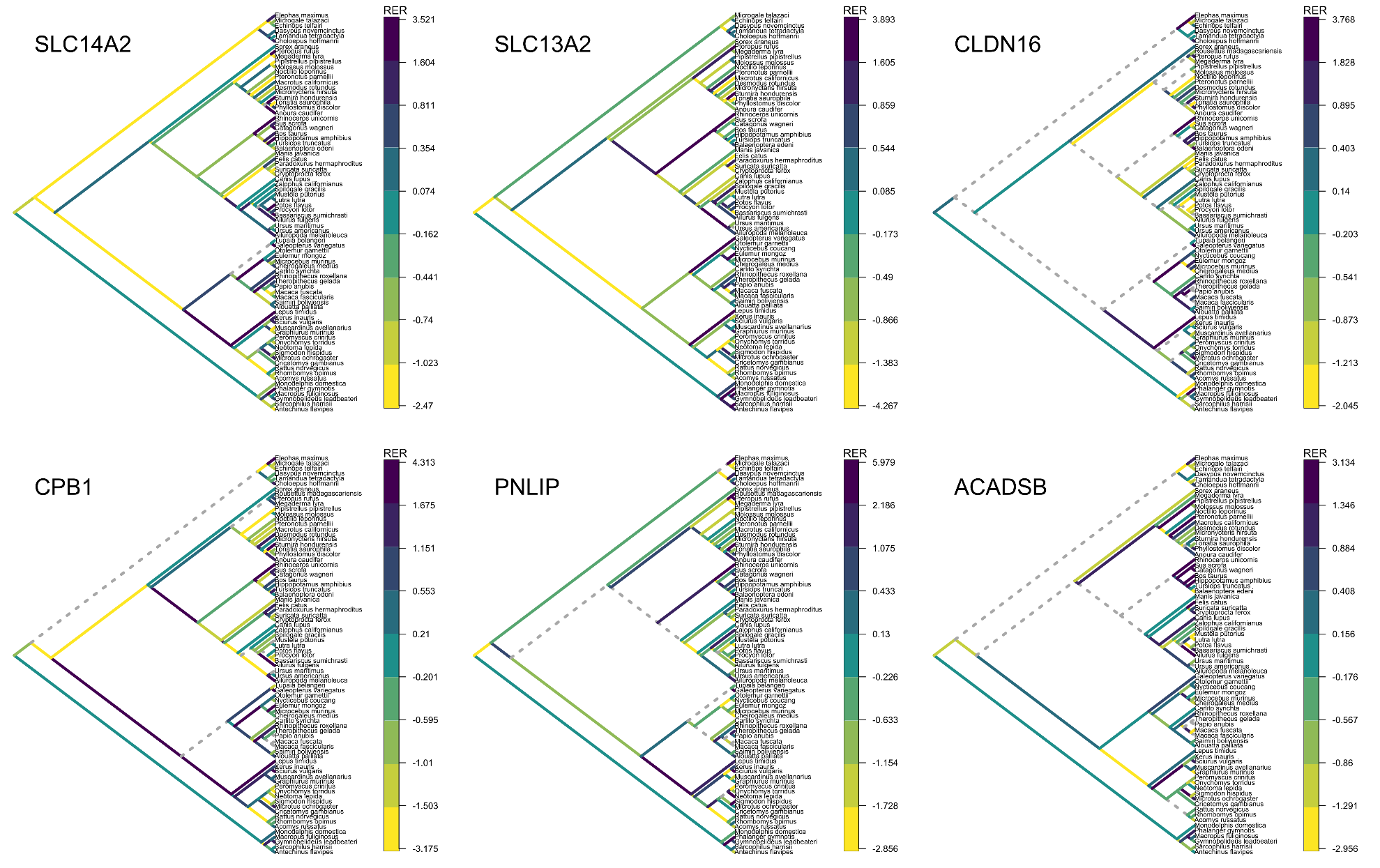

**Figure S1.** Relative evolutionary rates (RERs) across the mammalian phylogeny for genes significantly associated with change in carnivory score. Branch colors represent the RER for a given gene in each part of the phylogeny. Purple and yellow branches indicate positive and negative RERs, respectively. Gray dashed lines represent branches that were excluded from our analyses due to default branch length filtering in RERconverge, which set the shortest 5% of branches across all gene trees to N/A. In each gene, positive and negative RERs are distributed throughout the phylogeny.

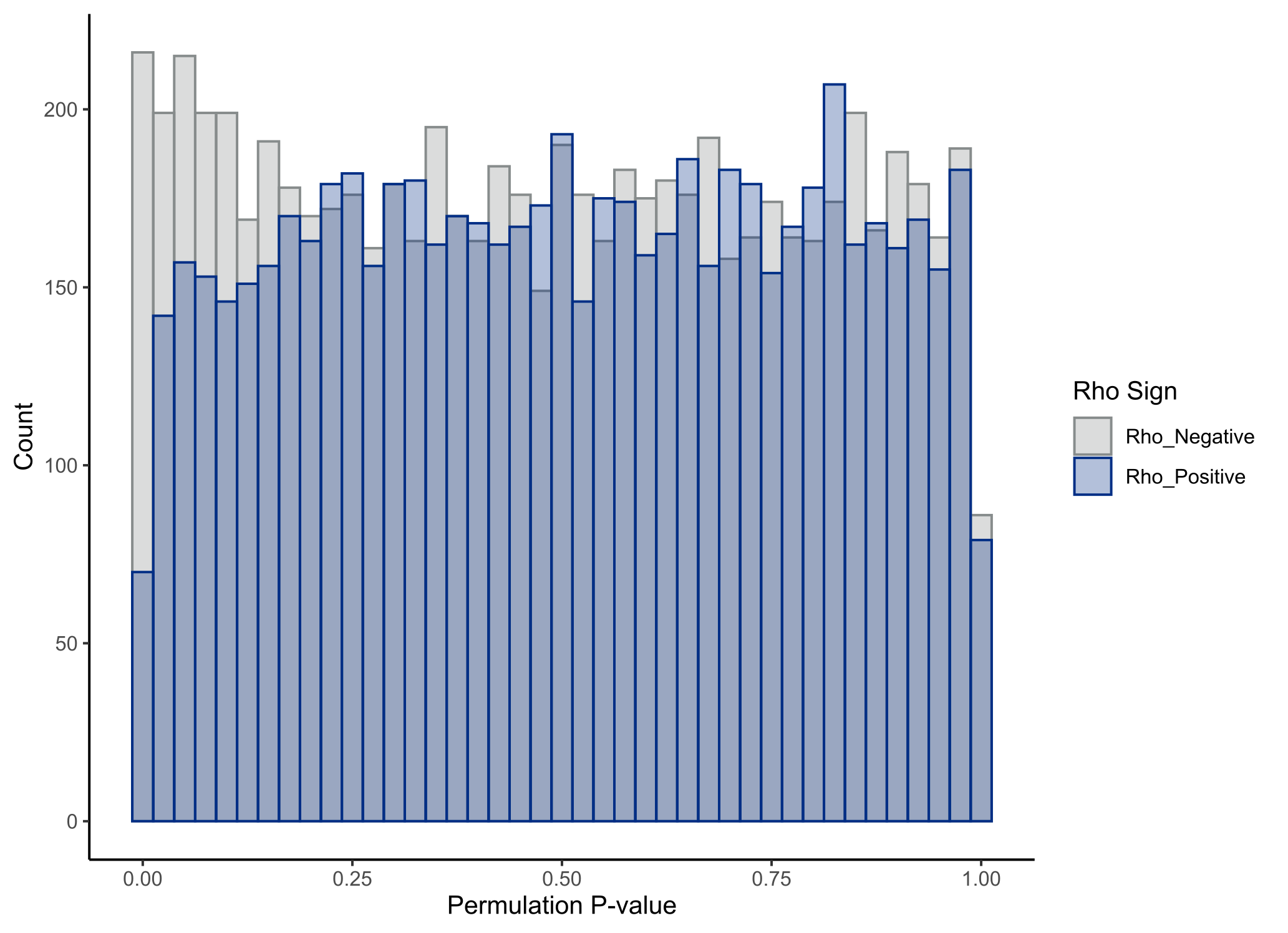

**Figure S2.** Significant differences in the distribution of empirical, permulation-based *P*-values for positive and negative associations between gene relative evolutionary rates (RERs) and carnivory score (Mann-Whitney U test, AUC=0.523, *P*=2.16 x 10^-6^). Our analysis showed reduced power to detect positive associations between RER and change in carnivory, as indicated by fewer low *P*-values being generated during permulation for genes with a positive correlation.

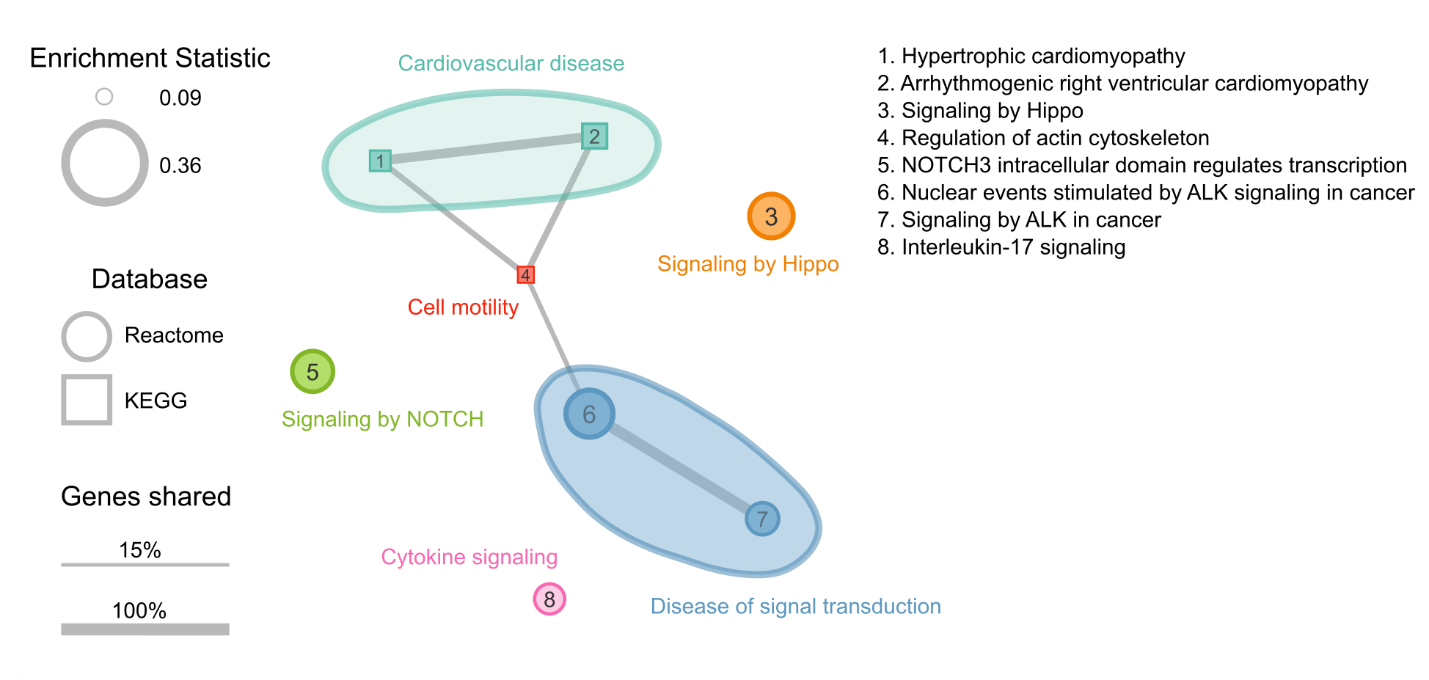

**Figure S3.** Significantly enriched gene pathways (n=8) under less evolutionary constraint as carnivory score increases. Each circle or square represents a gene pathway. Circles and squares represent pathways from the Reactome and KEGG databases, respectively. The size of the shape represents the magnitude of the difference between the distribution of test statistics for genes in that pathway and the distribution for all other genes. Larger shapes, representing pathways with larger positive correlation statistics, indicate greater reductions in evolutionary constraint as carnivory score increases. The width of lines connecting pathways represents the proportion of shared genes in the smaller gene set. Colors represent the broad functional categories that the pathways occupy.

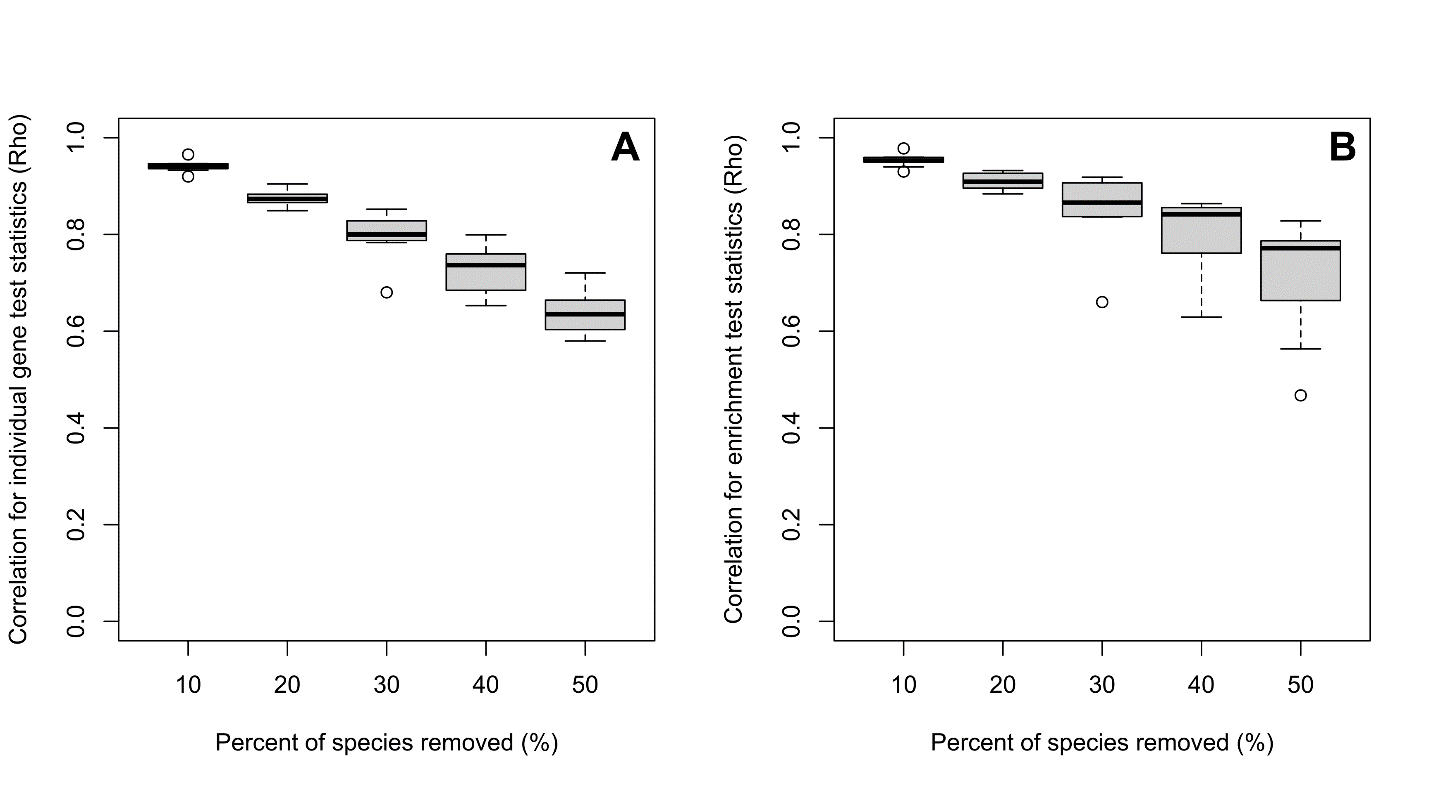

**Figure S4.** Correlations between RERconverge results using our full species list and using species subsets where 10, 20, 30, 40, or 50 percent of species have been randomly removed. For each percent of species removed, we created 10 random subsets. Boxes represent the distribution of correlation values (Pearson’s correlation coefficient, rho) between the original results and the results with a given percent of species removed. (*A*) Correlations between gene correlation statistics. (*B*) Correlations between gene set enrichment statistics.

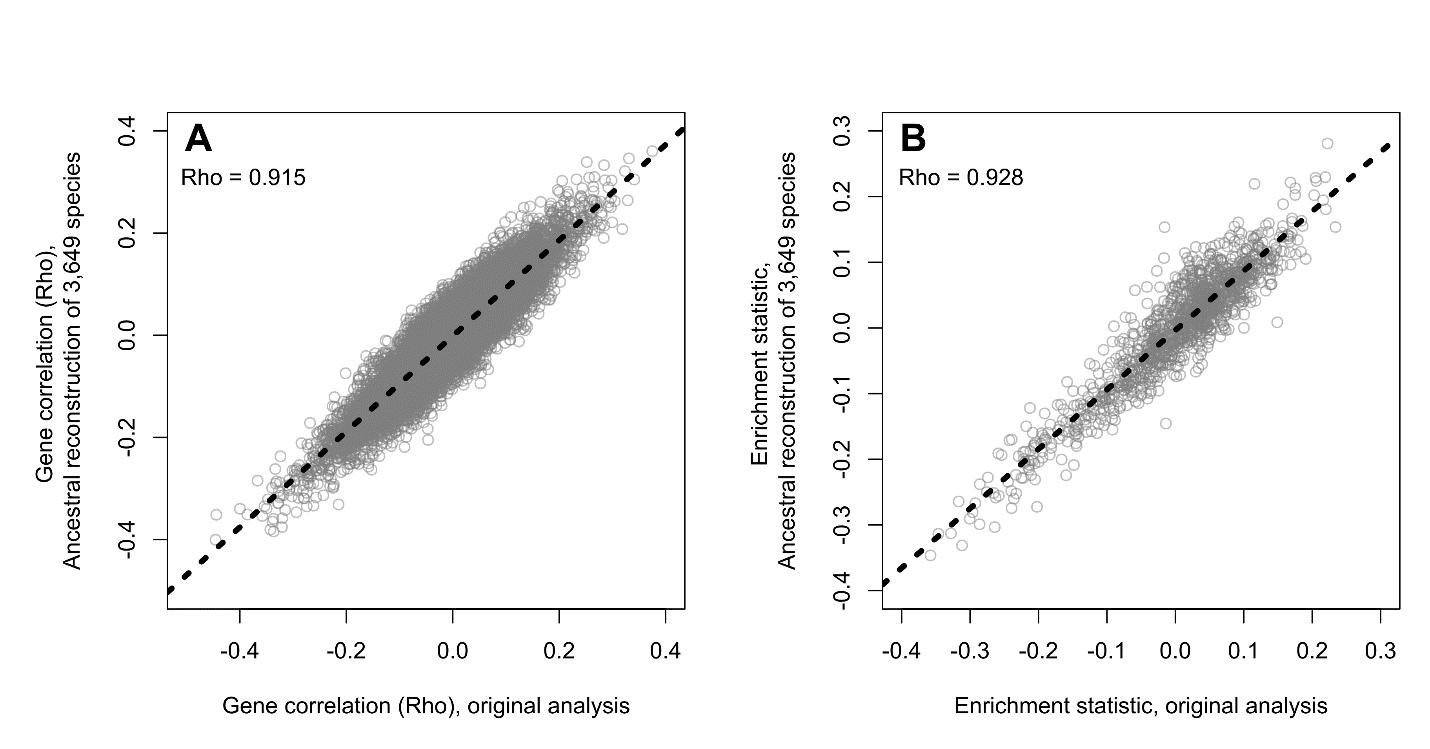

**Figure S5.** Correlations between RERconverge results using the original ancestral state reconstruction method built into RERconverge and a modified reconstruction method. For the modified method, we performed fast estimation of maximum likelihood ancestral states (Revell 2012) on a larger tree of 3,649 species for which we could obtain diet data from EltonTraits (Wilman et al. 2014). We then pruned this reconstruction to our study species and used it as input in the RERconverge analysis. Correlations were calculated using Pearson’s correlation coefficient (rho). Dashed lines indicate the regression line. (*A*) Correlation between gene correlation statistics. (*B*) Correlation between gene set enrichment statistics.

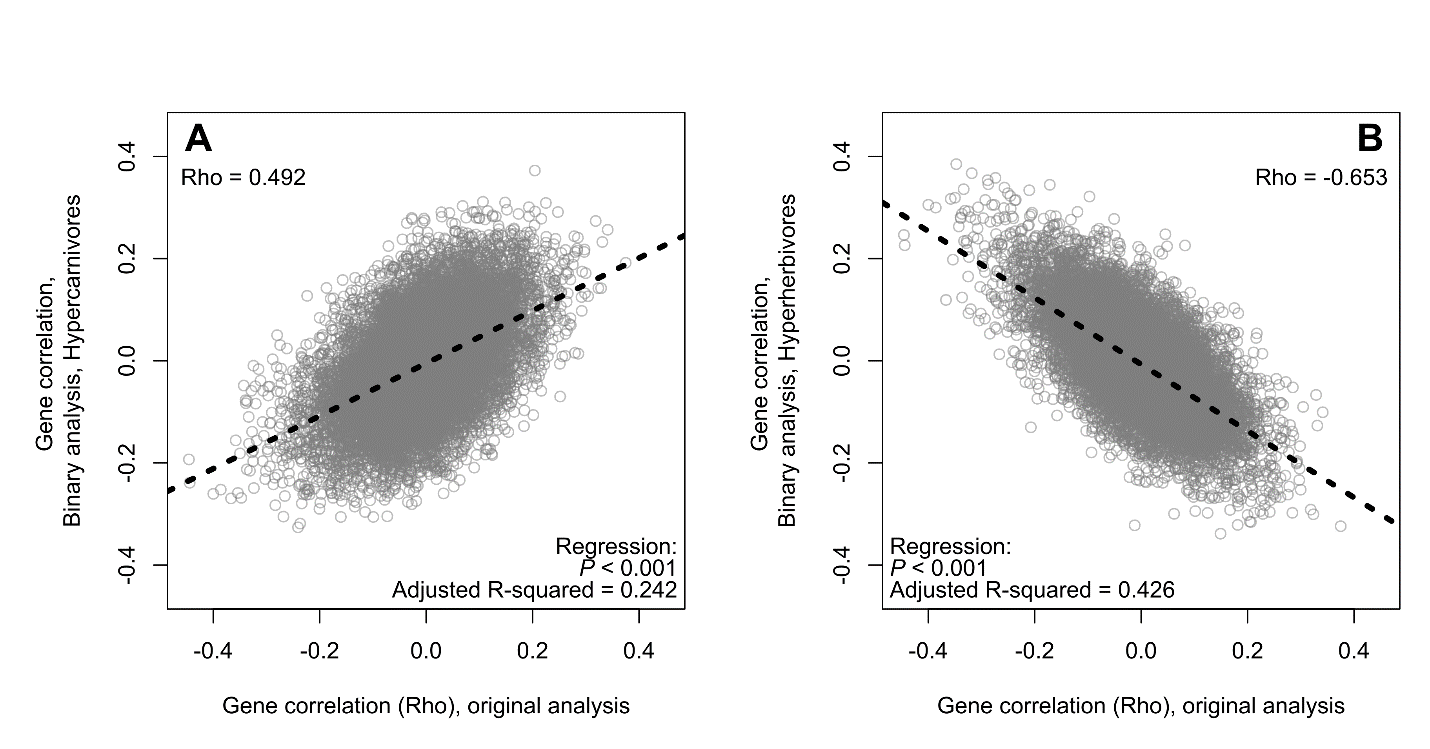

**Figure S6.** Correlations between the test statistics for the continuous and binary RERconverge analyses. Inset rho values represent the Pearson correlation coefficient between the two analyses. Dashed lines indicate the regression line. (*A*) Correlation between the individual gene test statistics from the original continuous analysis and the binary analysis with hypercarnivores (carnivory scores ≥90) as foreground. (*B*) Correlation between the individual gene test statistics from the original continuous analysis and the binary analysis with hyperherbivores (carnivory scores ≤10) as foreground.

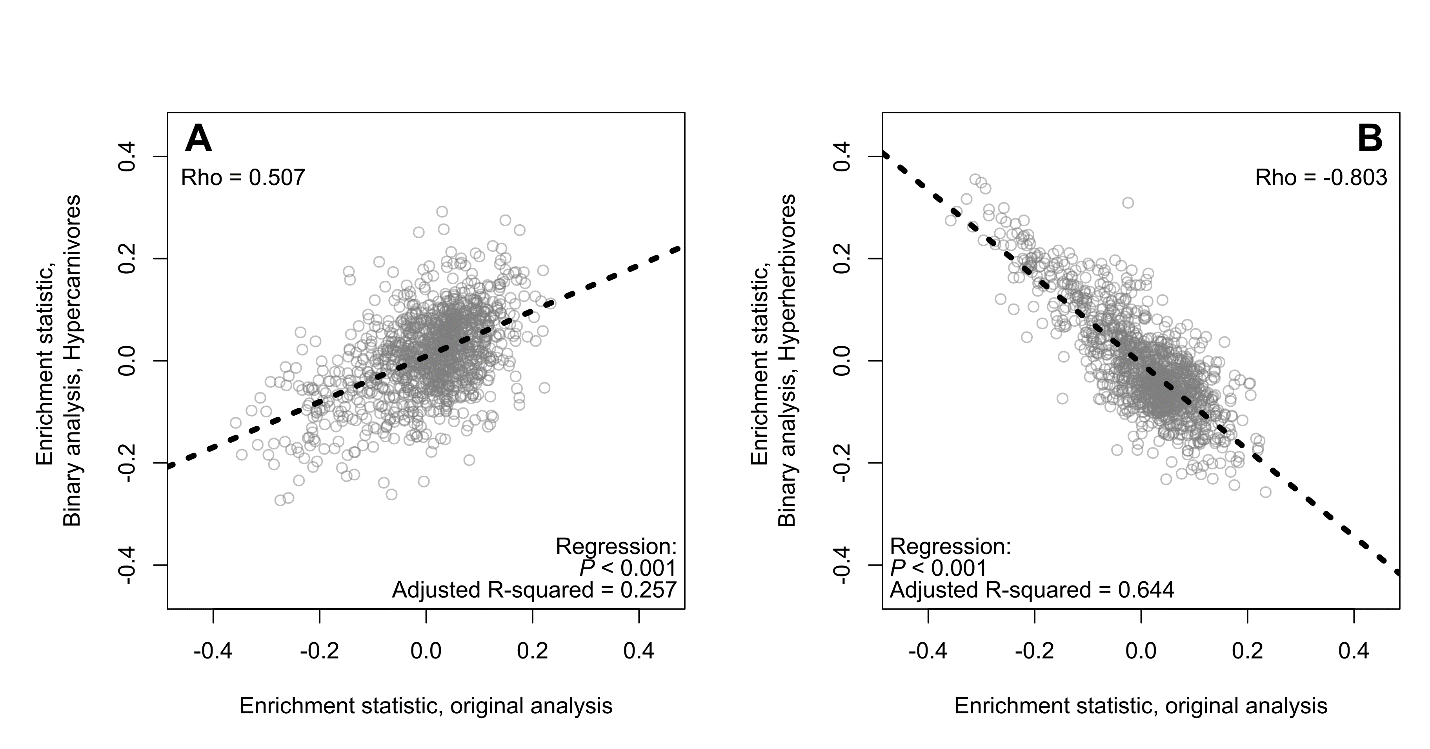

**Figure S7.** Correlations between the gene pathway enrichment statistics for the continuous and binary RERconverge analyses. Inset rho values represent the Pearson correlation coefficient between the two analyses. (*A*) Correlation between the enrichment statistics from the original continuous analysis and the binary analysis with hypercarnivores (carnivory scores ≥90) as foreground. (*B*) Correlation between the enrichment statistics from the original continuous analysis and the binary analysis with hyperherbivores (carnivory scores ≤10) as foreground.

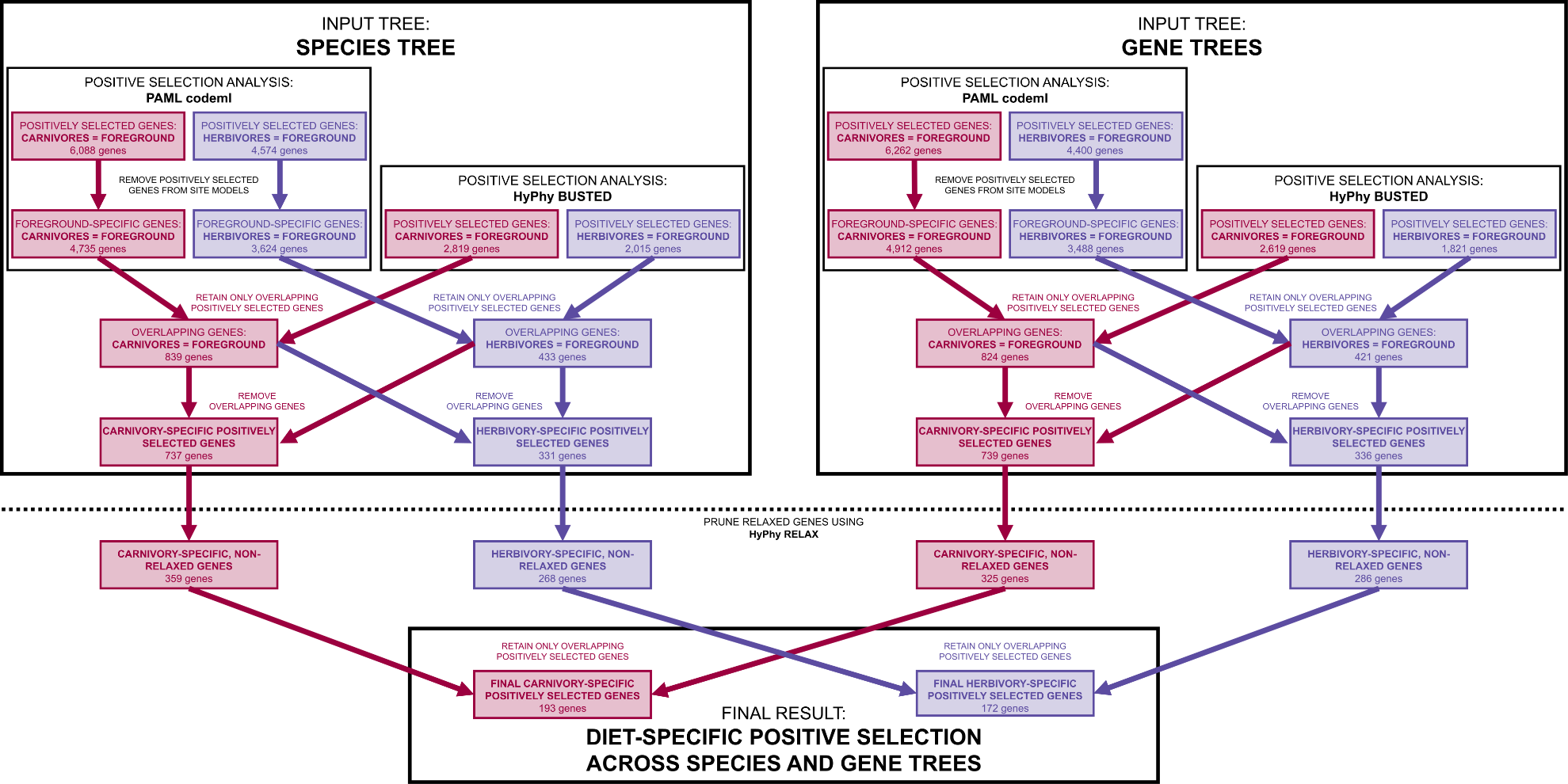

**Figure S8.** Conservative strategy for generating final lists of positively selected genes associated with carnivory and herbivory in mammals. We tested 15,117 genes for positive selection in carnivores (species with carnivory scores ≥90) and herbivores (species with carnivory scores ≤10) separately, using both codeml from PAML (Yang 2007) and BUSTED from HyPhy (Murrell et al. 2015; Pond et al. 2020). To account for the impact of gene tree discordance, we performed our positive selection analyses using both the species tree (Upham et al. 2019) and gene trees inferred using RAxML (Stamatakis 2014). We only considered a gene to have experienced diet-associated positive selection when it produced a significant result in both our codeml and BUSTED analyses, and if we got this result using both the species and gene trees. As an additional filtering step, we tested the genes returned by codeml and BUSTED for relaxed selection using RELAX (Wertheim et al. 2015) and excluded any genes showing significant relaxation in the foreground species from our final gene lists. The final products of this process were two conservative lists representing positively selected genes in the most carnivorous and herbivorous mammals, respectively (Tables S7, S8).
