## Supplemental_Methods for "Convergent relaxation of molecular constraint in mammalian herbivores highlights the roles of liver and kidney functions in carnivory"

**Supplemental Material**

**ADDITIONAL METHODS**

***Codon alignment generation and filtering***

The codon alignments used in our study were generated by the Zoonomia Consortium using TOGA and MACSE v2 (Ranwez et al. 2018; Kirilenko et al. 2023). The method used by the Consortium to generate the alignments is not detailed in the associated paper, but is described online at: <https://genome.senckenberg.de/download/TOGA/README.multipleCodonAlignments.txt> [last accessed: 2024/04/23]. For easy reference, we repeat their methodology below:

“*1) For each gene annotated in the reference species, we selected the transcript with the longest CDS.*

*2) For a query species, we considered orthologs for this transcript that were classified as Intact, Partial Intact or Uncertain Loss.*

*3) Two steps assure that alignments mostly (but not exclusively) include 1:1 orthologs. First, if a query assemblies has more than four predicted orthologs, we omitted this query assembly. Second, if the gene does not have a single (1:1) ortholog for at least 75% of all query species, we did not compute a multiple codon alignment for this gene.*

*Before running MACSE2.0, TOGA masked all inactivating mutations (frameshifting indels or premature stop codons) in all query sequences by replacing them with XXX codons. Each transcript was split into individual exons, and each exon was aligned individually with MACSE2.0. The exon alignments, with potential codons that are split between exon boundaries, were concatenated into a final alignment for the entire gene. This procedure ensures that multiple codon alignments are accurate and therefore suitable for selection screens*”.

We then employed the filtering methodology described in Wirthlin et al. (2024) to these alignments, before running additional filtering steps as described in our study. For easy reference, the filtering methodology of Wirthlin et al. (2024) is repeated below:

“*These alignments were subsequently filtered to remove duplicated species, poorly represented proteins, and low-scoring alignments. Specifically, alignments with fewer than 221 unique species (0.025 quantile of the distribution of unique species number for all alignments), alignments with fewer than 189 total species with ungapped coverage of 50% of the total alignment length (0.1 quantile), alignments with more than 97 duplicated species (0.95 quantile), and alignments with ungapped length <267 bp (0.01 quantile) were excluded. In total, this resulted in excluding 4,723 transcripts representing 2,613 unique genes. Within the remaining alignments, any sequences that did not cover 50% of the total alignment length were excluded, and, when there were multiple sequences for a species within an alignment, the sequence with the highest identity to the human reference sequence across the full alignment length was retained.*”

***Robustness of RERconverge results***

We assessed the sensitivity of our RERconverge results to the choice of species included in our analysis. We created subsets of our species list by randomly removing 10, 20, 30, 40, or 50 percent of species. For each percent of species removed, we created 10 random subsets. We then ran the RERconverge analysis with these subsets, without permulations. We also tested for enrichment of KEGG and Reactome pathways. To quantify how robust our results are to species selection, we calculated the correlations between our full species analysis and the subset analyses for both gene correlation and pathway enrichment statistics.

We assessed the sensitivity of our RERconverge results to changes in the ancestral state reconstruction of the carnivory score. The standard ancestral reconstruction performed with RERconverge is limited to the species included in the analysis, so it does not account for diet evolution that occurred outside of our lineages of interest. This may make inferences of the timing, magnitude, and direction of diet evolution less accurate. We performed fast estimation of maximum likelihood ancestral states (Revell 2012) on a larger tree of 3,649 species for which we could obtain diet data from EltonTraits (Wilman et al. 2014). We pruned this reconstruction to our study species and used it as input in the RERconverge analysis, without permulations. We then tested for enrichment of KEGG and Reactome pathways. To quantify how robust our results are to ancestral diet reconstruction methods, we calculated the correlations between our original results and those of the new reconstruction method for both gene correlation and pathway enrichment statistics.

***Binary RERconverge analyses***

In addition to our main RERconverge analysis using a continuous carnivory score, we performed analyses of relative evolutionary rate (RER) using binary classifications of diet. We used the binary version of RERconverge to find genes with significant increases or decreases in RER in the most carnivorous or herbivorous lineages. We used the same species in our carnivore and herbivore foregrounds as in our analyses of positive selection (Fig 1; see Methods) but used the ‘clade = all’ option, wherein all lineages following a major phenotype transition are considered foreground. We also allowed bidirectional transitions between foreground and background to occur along the phylogeny. We performed 10,000 permulations of each analysis using the *getPermsBinaryFudged* function and corrected for multiple hypothesis testing using Storey’s correction method (Storey et al. 2020). The *getPermsBinaryFudged* function allows a permulated tree to differ from the original in the number of total foreground species, up to a value specified by the ‘fudge’ parameter. We used the default ‘fudge’ value of five. We revised the permulation functions to enable bidirectional transitions between foreground and background within the permulated phenotype trees. These updates are currently available in the ‘AddBidirectionalPerms’ branch of the RERconverge repository on GitHub (<https://github.com/nclark-lab/RERconverge/tree/AddBidirectionalPerms>).
